# Multi-center integrated analysis of non-coding CRISPR screens

**DOI:** 10.1101/2022.12.21.520137

**Authors:** David Yao, Josh Tycko, Jin Woo Oh, Lexi R. Bounds, Sager J. Gosai, Lazaros Lataniotis, Ava Mackay-Smith, Benjamin R. Doughty, Idan Gabdank, Henri Schmidt, Ingrid Youngworth, Kalina Andreeva, Xingjie Ren, Alejandro Barrera, Yunhai Luo, Keith Siklenka, Galip Gürkan Yardımcı, The ENCODE4 Consortium, Ryan Tewhey, Anshul Kundaje, William J. Greenleaf, Pardis C. Sabeti, Christina Leslie, Yuri Pritykin, Jill E. Moore, Michael A. Beer, Charles A. Gersbach, Timothy E. Reddy, Yin Shen, Jesse M. Engreitz, Michael C. Bassik, Steven K. Reilly

## Abstract

The ENCODE Consortium’s efforts to annotate non-coding, *cis*-regulatory elements (CREs) have advanced our understanding of gene regulatory landscapes which play a major role in health and disease. Pooled, non-coding CRISPR screens are a promising approach for systematically investigating gene regulatory mechanisms. Here, the ENCODE Functional Characterization Centers report 109 screens comprising 346,970 individual perturbations across 13.3Mb of the genome, using a variety of methods, readouts, and statistical analyses. Across 332 functionally confirmed CRE-gene links, we identify principles for screening endogenous, non-coding elements for causal regulatory mechanisms. Nearly all CREs show strong evidence of open chromatin, and targeting accessibility peak summits is a critical component of our proposed sgRNA design rules. We provide experimental guidelines to accurately detect CREs with variable, often low, transcriptional effects. We discover a previously undescribed DNA strand-bias for CRISPRi in transcribed regions with implications for screen design and analysis. Benchmarking five screen analysis tools, we find CASA produces the most conservative CRE calls and is robust to artifacts of low-specificity sgRNAs. Together, we provide an accessible data resource, predesigned sgRNAs targeting 3,275,697 ENCODE SCREEN candidate CREs, and screening guidelines to accelerate functional characterization of the non-coding genome.

## Introduction

The non-coding genome contains critical regulators of gene expression and harbors >90% of trait-associated human genetic variation^1–4^. Major efforts over the past decade have mapped hundreds of thousands of non-coding, candidate *cis*-regulatory elements (cCREs)^5–7^. Such efforts have relied primarily on mapping sequence conservation and biochemical markers that are correlated with regulatory activity, rather than direct functional characterization. Site-specific, programmable, and highly scalable CRISPR genome and epigenome manipulation methods have enabled massively parallel perturbation assays to identify and characterize functional *cis*-regulatory elements (CREs). In this text, CREs are the elements with empirically characterized endogenous function, whereas cCREs refer to uncharacterized elements–the targets of a screen, often nominated by biochemical markers– or those which did not have significant effects in a screen but may still have regulatory function in another biological context.

Various CRISPR-based perturbation methods have been developed to determine a cCRE’s effects on target gene expression and/or downstream phenotypes^8–14^. To date, systematic and comprehensive benchmarking of non-coding CRISPR screening methods, and attempts to harmonize the results, have been limited by the low number of available datasets and inconsistent reporting. We reasoned that leveraging our large resource of datasets with varied library design approaches, perturbation modalities, phenotypic readouts, and analysis methods, would enable us to distill useful guidelines for successful execution and analysis of these complex experiments. This effort would also provide a uniquely robust and well-suited dataset for generating and rigorously testing hypotheses about gene regulation that would otherwise be difficult to assemble as a single laboratory.

The ENCODE4 Functional Characterization Centers have generated the largest collective dataset of endogenous cCRE perturbation screens to date, using diverse experimental approaches. Here we compare non-coding CRISPR screening approaches, and propose standardized practices and data file formats generalizable to all such screens. We highlight how particular methodologies and experimental parameters can be tuned to address specific biological questions and technical limitations. By assembling and jointly analyzing this large repository of bulk CRISPR screens, we develop suggested practices for study design, analysis, and validation, as well as provide comprehensive benchmarking between methodologies. We analyze the strengths and weaknesses of various CRISPR non-coding screens at each screening stage including: 1) library design, 2) CRISPR perturbation selection, 3) phenotyping strategy, and 4) analytical methods. Finally, we leverage our combined analysis of >100 distinct CRISPR screens to interrogate broader properties of gene regulation.

## Results

### Diverse approaches of the ENCODE non-coding CRISPR database identify functional regulatory elements overlapping markers of CREs

The ENCODE4 Functional Characterization Centers have undertaken the largest set of diverse non-coding CRISPR screens to date, performing >100 experiments in human and mouse biosamples, all of which are available on the ENCODE portal^15^ (see **Supplementary Section 1**). The data used in this study are divided into three targeting approaches (**Fig. 1A, Supplementary Tables 1-3**): 1) unbiased tiling screens that include sgRNAs targeting cCREs and non-cCRE regions within a specific locus (*e*.*g*., an entire topologically associated domain (TAD))^9,10,16^, 2) screens that select sgRNAs targeting cCREs in a given locus^12,17,18^ and 3) screens that target cCREs in multiple loci or genome-wide^19^. The different approaches have different strengths. For example, tiling screens can identify novel CREs that lack epigenetic marks commonly associated with regulatory activity, while cCRE-targeted approaches can screen many more putative regulatory elements with the same number of sgRNAs.

**Fig. 1.**
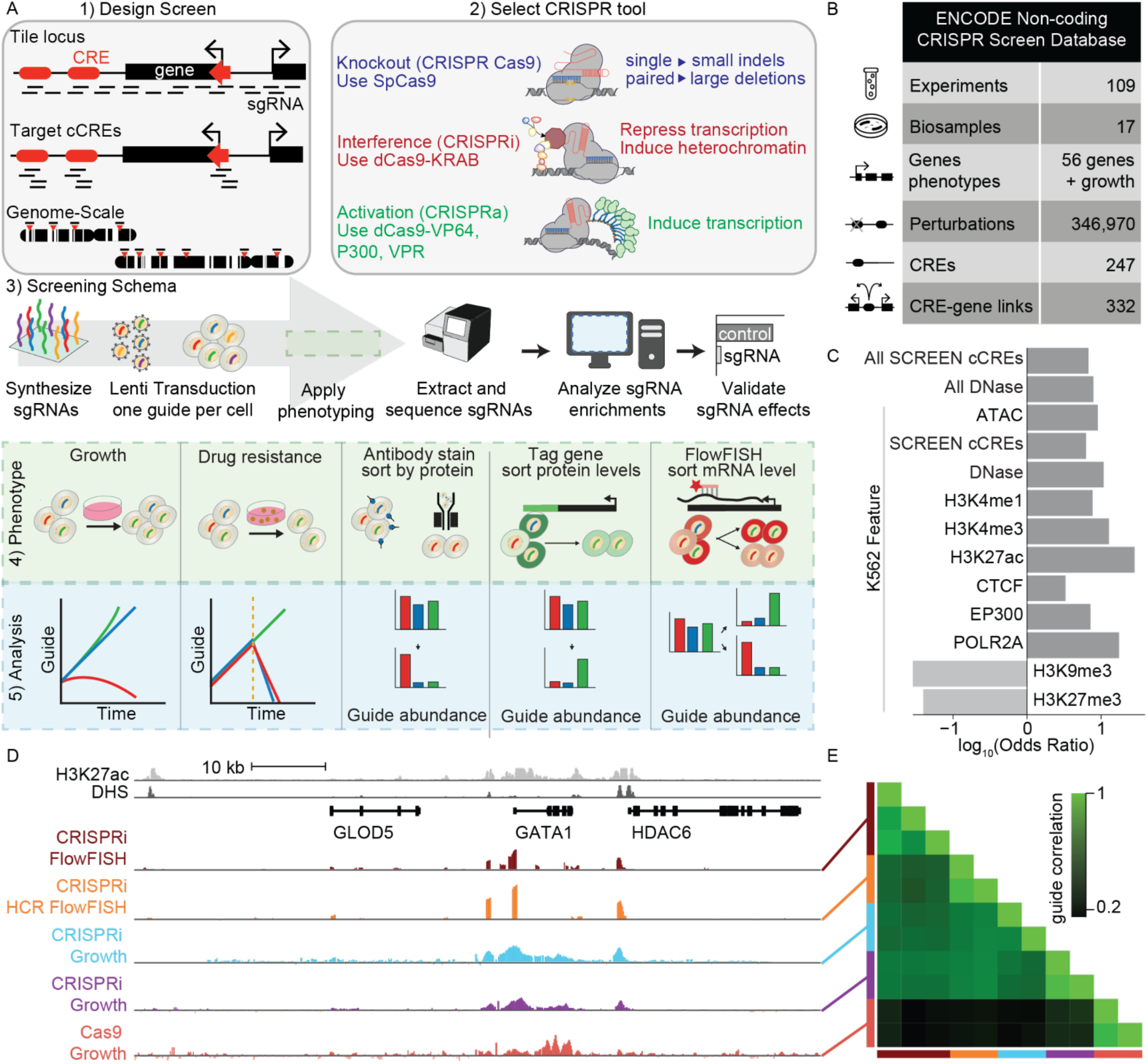
The ENCODE non-coding CRISPR Screening Database. **A**) Overview of CRISPR non-coding strategies including 1) perturbation design strategies, 2) CRISPR perturbation modalities, 3) workflow of a standard screen, 4) phenotyping strategies, and 5) analysis approaches. **B**) CRISPR screen data summary from the April 2022 release of the ENCODE portal. “Experiments”, “Biosamples”, and “Genes/phenotypes” reflect all human CRISPR screens. “Perturbations”, “CREs”, and “CRE-gene links” reflect results of K562-focused analysis. **C**) Odds ratio for genomic annotation overlap with CRISPR screen-identified regulatory elements (N=210, **Methods**). “All” refers to cell-agnostic features. K562 refers to cell-type annotations. All odds ratios were significant at P value<0.01 and values were log_10_-transformed for visualization (two-sided Fisher’s exact test). **D**) Genome browser snapshot of the *GATA1* locus including H3K27ac (light gray) and DHS signal (dark gray) in K562 cells. CRISPR screen data (signal log_2_FC) for one replicate each of CRISPRi-FlowFISH (dark red), CRISPRi-HCR-FlowFISH (orange), Tycko et. al. 2019 CRISPRi-Growth (blue), Fulco et. al. 2019 CRISPRi-Growth (purple), and Cas9-Growth (red). **E**) Pairwise correlations (Pearson) of normalized effect sizes on sgRNAs commonly shared by each method (N=176 sgRNAs with GuideScan-aggregated CFD specificity score>0.2) in *GATA1* locus; multiple rows for the same CRISPR method show replicates.

Across each targeting strategy, three major CRISPR perturbation strategies were used: 1) genetic perturbations, including creation of small insertions and deletions induced by Cas9 nuclease (Cas9)^20,21^ and paired sgRNA deletions using Cas9 to excise large genomic regions (∼2-20 kb)^8,16,22^; 2) epigenetic repression, with deactivated Cas9 (dCas9) fused to a KRAB domain to target transcriptional repression (CRISPRi)^23–25^; or 3) transcriptional activation, with dCas9 fused to activator domains (CRISPRa)^26–28^ (**Fig. 1A**). Following targeting design and perturbation choice, all screens introduced sgRNAs into cells at low multiplicity of infection (MOI) via lentiviral transduction. Next, a bulk phenotyping selection method was applied^9–12,14,16–18,22,29–31^, sgRNAs were sequenced, and differences in sgRNA abundance were quantified correlating with the measured phenotype. The ENCODE CRISPR screening database contains over 650,000 individual perturbations covering 13.3 Mb (0.43%) of the human genome (**Methods**). Regulatory activity was assayed for 56 genes and growth-related phenotypes in 19 human cell lines or induced pluripotent stem cells, collectively identifying 887 regions that, when perturbed, significantly impacted a cellular phenotype (**Supplementary Tables 1-2, Methods**).

Of the human cell lines, the most experiments were performed in K562 cells and, thus, we leveraged data from 53 distinct non-coding CRISPR screens with various experimental designs to gain insights into the characteristics and features that define CREs. We first performed a meta-analysis integrating data from these experiments and found that 230.6 kb (2.94%) of the 7.8 Mb perturbed in ≥1 experiment displayed control of gene expression or cellular growth (**Fig. 1B, Supplementary Table 1, Methods**). To identify the types of genomic states that best predict regulatory activity, we intersected the identified regions, referred to as CREs (n=210, **Supplementary Table 4**), with annotations of K562 cells and observed the greatest overlap with ENCODE SCREEN cCREs (88.6%, 186/210; Fisher’s exact test, P=4.93E-24, OR=6.27), and H3K27ac, RNA Pol-II, and H3K4me3 peaks, having the greatest enrichment (OR=28.3, 17.5, 12.8 respectively, P<1e-5 for each; **Fig. 1C, Supplementary Tables 5-6**). Similar enrichments were observed for the union set of ENCODE SCREEN cCREs and DNase hypersensitive sites (DHSs) annotated in all ENCODE biosamples (**Supplementary Fig. 1A, Supplementary Table 6**). Together, these results suggest that many commonly associated features of regulatory activity in genomics studies are also largely indicative of regulatory activity in non-coding CRISPR screens.

We questioned which genomic and biochemical features characterized the CREs identified in these non-coding CRISPR screens. The vast majority of CREs overlapped either accessible chromatin regions or H3K27ac peaks (95.7%, 201/210, **Supplementary Fig. 1B**). However, twenty-five CREs are marked by H3K27ac peaks but do not overlap DHSs, and 23 overlap DHSs but lack H3K27ac peaks (11.9% and 11.0% respectively). In K562 cells specifically, 9 CREs lack either of these features, but 7 of those 9 elements are located within DHSs in at least 1 other ENCODE biosample. Of the two remaining elements without overlap of any feature, one is a distal intergenic region ∼7.7kb upstream of the gene *EXOC1L*, and directly overlaps an annotated NFIC ChIP-seq peak in K562 cells. Similarly, the second element is located within an intron of the gene *FADS2* and is within 100 bp of an RBFOX2 ChIP-seq peak in K562 cells. Given these transcription factors (TFs) are implicated in cancer and cellular proliferation^32,33^, these results suggest the corresponding phenotypic changes may be driven by disruption of transcription factor-mediated regulatory interactions.

We next compared the quantitative signal of various accessibility measures, histone modifications, and *trans*-factor occupancies to interrogate which feature(s) best defined CREs identified in CRISPR screens. We observed that CREs have a greater mean signal than perturbed, non-significant regions, for chromatin accessibility, CTCF, EP300, H3K4me3, H3K27ac, and RNA Pol II binding, a lesser mean signal for H3K4me1 and H3K27me3, and no significant difference in H3K9me3 signal (**Supplementary Fig. 1C, Supplementary Table 7**). Overall, these results suggest that overlap and signal strength of accessible chromatin and active histone marks identify the majority of functional regulatory elements. However, we observed that some regulatory elements exhibit different combinations of epigenomic features (**Supplementary Fig. 1B**), in agreement with previous functional characterization experiments performed outside of ENCODE in which enhancers identified using MPRA could be classified into subclasses based on epigenetic features and reporter activity^34^.

### Individual sgRNA validations and effects of common sgRNAs demonstrate CRISPR screen results are reproducible

To examine the reliability of the datasets, we assessed the agreement between CRE effect sizes and individual sgRNA validations from a subset of experiments performed in K562 cells across multiple phenotypic readouts and analysis methods^9,10,12,17,35^. We compared the fold change in gene expression, measured via RT-qPCR, to the enrichment or depletion of individual sgRNAs in FACS-based (2 experiments, 2 genes) and growth-based CRISPR screens (2 experiments, 2 genes) (**Supplementary Fig. 2A-D**). We found that the screen result, computed on an element-level (**Supplementary Fig. 2A, B**) or on an individual sgRNA-level (**Supplementary Fig. 2C, D**), significantly correlates with the change in mRNA expression of the CRE’s target gene in individual sgRNA validation experiments (R^2^>0.75 for all screens). These individual sgRNA results confirm the quality of screen datasets where validation data was available. Validation methods to confirm CRE activity are further described in **Supplementary Section 2**.

To interrogate how different screening approaches compared at the same CREs, we identified sgRNAs used multiple times across sixteen screens at two commonly studied loci, *GATA1* (**Fig. 1D**) and *MYC* (**Supplementary Fig. 3A**). The underlying library size and targeting overlap of the *GATA1* and *MYC* screens varied (**Fig. 1D, Supplementary Fig. 3B, C**). Altogether, these screens deployed over 140,000 individual sgRNA, perturbing 1,655 cCREs in *GATA1* and *MYC* flanking regions that are 1.3 Mb and 2.0 Mb wide, respectively. For the 176 sgRNAs common between all five *GATA1* screens (after filtering with GuideScan^36,37^ CFD specificity scores≥0.2 to reduce possibly confounding off-target effects^17^), we observe strong replication within a screening approach (n=5; Pearson Correlation; min: 0.59, max: 0.90, mean: 0.77). For CRISPRi, there was strong correlation between experiments (n=36; Pearson Correlation min: 0.42, max: 0.90, mean: 0.56). In contrast, there was low (n=18; Pearson Correlation min: 0.15, max: 0.32, mean: 0.21) correlation between CRISPRi and Cas9 tiling experiments (**Fig. 1E**). CRISPRi experiments, whether interrogated by HCR-FlowFISH or growth, identified similar CREs at the MYC locus as well (**Supplementary Fig. 3A**). We reason it is because small indel mutations induced by Cas9 elicit greater effects in exons by causing nonsense mediated decay, whereas CRISPRi elicits greater effects in promoters and enhancers by perturbing essential mechanisms of gene regulation (**Supplementary Fig. 3D**).

### Integrated analysis of CRISPR screens provides guidelines for selecting cCRE targets and sgRNAs

We next sought to improve the selection of sgRNAs for non-coding CRISPRi screens and determine how to balance scale, sensitivity, and practicality. To do so, we analyzed 15 highly sensitive CRISPRi HCR-FlowFISH screens perturbing over 100 kb at 8 loci in K562 cells^8–10,16^. While dCas9 and Cas9 can be used to interrogate cCRE function, they are not ideal for discovering CREs in non-tiling screens due to their limited perturbation ranges and effect sizes, and are thus excluded from this discussion.

The HCR-FlowFISH datasets were comprehensive tiling screens that targeted all regions regardless of cCRE or other epigenetic feature prioritization; however, for cCRE-targeting approaches, defined targets can be prioritized based on epigenetic features. Consistent with our findings above, we observed that the significant CREs identified across the HCR-FlowFISH screens are enriched in accessible chromatin (74%) or H3K27ac ChIP-seq peaks (80%; 87% have at least one feature), with the overwhelming majority having both epigenetic features (**Supplementary Fig. 4A**). Thus, a combination of these and other CRE-associated epigenetic features (**Supplementary Fig. 1B**) can be used to nominate cCRE targets. As enhancers can be thousands to millions of nucleotides away from their target gene, screening all potential cCREs in this range may not be feasible^12,38^. In such cases, prioritizing cCREs <100 kb away from target genes was sufficient to identify 90% of the significant CREs, consistent with a previous single-cell study^39^. cCREs can be selected within topologically associating domains (TADs), and all significant CREs from the HCR-FlowFISH screens were found to be within the same TAD as their target gene^8–10,16^. Predictive modeling using the Activity-by-Contact Model (ABC)^12,38^ identified 43% of the significant elements and contact-mapping datasets generated via promoter-capture Hi-C^40^ can also be used to prioritize cCREs–noting that some elements may be missed. Optimizing the sgRNAs targeting each cCRE is crucial for maximizing perturbation strengths without compromising practicality or scale. cCREs are often nominated based on DHS or H3K27ac peaks, which span hundreds of base pairs and therefore may provide flexibility in sgRNA choice and positioning. We compared sgRNA perturbation effects within significant, non-promoter enhancers and observed sgRNAs overlapping the DHS peak induced significantly stronger perturbations than those overlapping the H3K27ac peak (**Fig. 2A**, binomial test P value<0.001). Further, sgRNA effects across tiled loci revealed local perturbation maxima near the enhancer’s DHS summit (**Supplementary Fig. 4B**). Aggregating all significant enhancers together, we found that sgRNA effects are generally strongest nearest the DHS summit, with a near-linear decrease as a function of distance from the summit (**Fig. 2B, Supplementary Fig. 4C-D**).

**Fig. 2.**
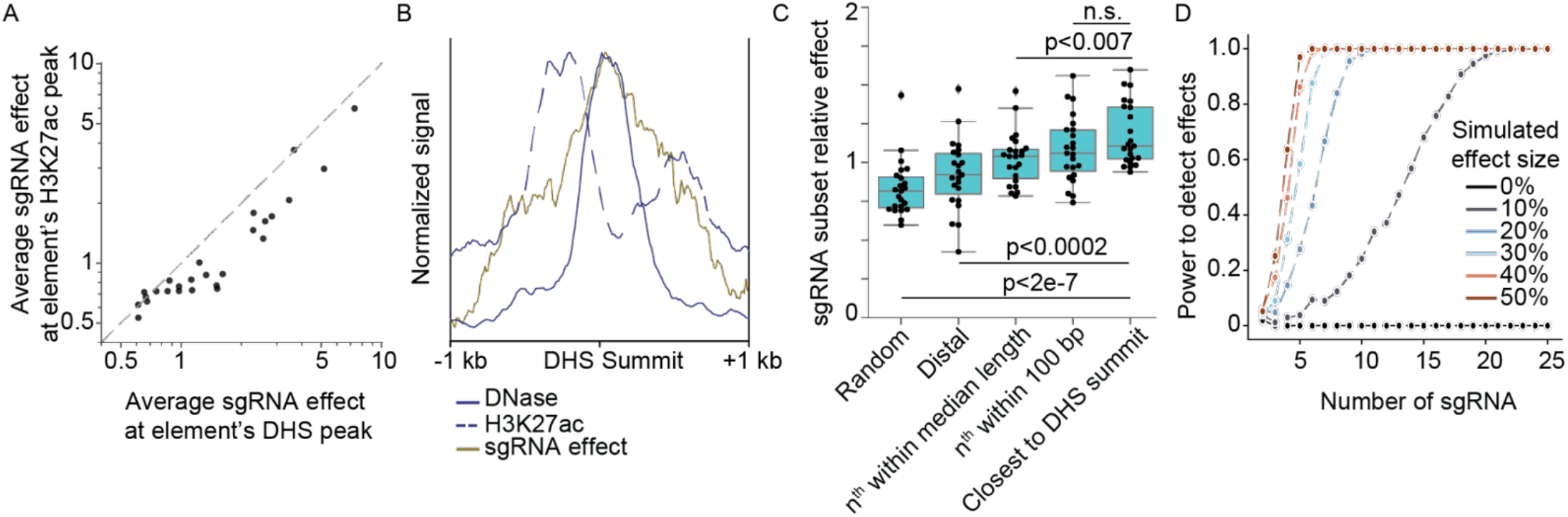
Integrated analysis of non-coding CRISPR screens provides guidelines for selecting cCRE targets and sgRNAs. **A**) Average effects of all sgRNAs within DHS or H3K27ac peaks at significant enhancers intersecting both epigenetic features. **B**) bigWig P value signal tracks for H3K27ac ChIP-seq and DNase-seq, and base pair-normalized effects of 6338 sgRNAs within +/- 1 Kb of the DHS summit, for 27 significant enhancers intersecting 32 DHS and H3K27ac peaks. **C**) Comparison of sgRNA selection strategies. Points reflect the effects of 10 sgRNAs selected by the indicated method for significant enhancers, normalized to the mean effect of all sgRNAs in that enhancer. ‘Random’ is the average of 100 random subsets from across the DHS peak. ‘Distal’ are sgRNAs closest to half the median DHS peak length (179 bp) from the summit. Every ‘n^th^’ sgRNA is selected by arranging sgRNAs in order of their PAM’s genomic coordinate, and selecting every n^th^ sgRNA such that their ranked orders are evenly spaced. ‘Closest’ sgRNAs are nearest to the DHS summit. Boxes show the quartiles, with a line at the median, lines extend to 1.5 times the interquartile range, and dots beyond lines show outliers. **D**) Power simulation to detect significant effects on GATA1 expression as a function of the element’s effect size and the number of sgRNAs. Power was computed by simulations using three replicates of GATA1 CRISPRi-FlowFISH data, where sgRNA effects in the eGATA1 element were scaled such that the average adjusted effect of all sgRNAs in the enhancer was 10-50%.

We compared methods for selecting sgRNA subsets and concordantly found that choosing sgRNAs closest to the DHS summit performed better than selecting sgRNAs further from the summit, randomly, or when evenly spaced apart (**Fig. 2C**). This selection method is straightforward, only requiring summit calls, which are standard output from peak callers such as MACS2^41^.

Next, we investigated the minimally sufficient number of sgRNAs needed to test a target’s significance at a given effect size. We performed a conservative power analysis on a *GATA1* FlowFISH screen^10^ and observed that 10 sgRNAs, selected randomly from within the significant *GATA1* enhancer, are required to provide over 95% power to detect enhancers with a 20% or greater effect on gene expression (**Fig. 2D**). Since this *GATA1* screen was well-powered and reproducible, 10 sgRNAs should be considered the minimum if coverage and replicates cannot be maintained or experimental noise is anticipated to be greater (**Supplementary Fig. 5A-B**).

sgRNA design parameters including specificity and sequence filters can strongly impact CRISPR screen results^17^. In evaluating these filters in ENCODE screens, we found that the magnitude of the effect differed between gene expression versus proliferation-based phenotypic screens. Low specificity sgRNAs often confound proliferation-based screens due to off-target toxicity^17^. While a GuideScan-aggregated CFD specificity score≥0.2 can filter them out, this removes a large proportion of sgRNAs that are near the DHS summits due to increased local GC content and Cas9 PAM availability (**Supplementary Fig. 5C**)^42^. However, we found that screens with gene expression readouts (e.g. HCR-FlowFISH) were not as sensitive to these off-target effects (**Supplementary Fig. 5D**, Odds ratio=1.432 vs 1.811 for *GATA1* HCR-FlowFISH vs. proliferation screens; Fisher’s exact test P value>0.2) — presumably because any off-target effects of such sgRNAs are less likely to impact the expression of a specific gene than a more general phenotype like cellular proliferation. Therefore, specificity filters as stringent as a GuideScan-aggregated CFD specificity score ≥0.2 may not be needed for HCR-FlowFISH. sgRNA spacer sequence also affects efficacy: sgRNAs containing the U6 promoter termination sequence (‘TTTT’)^43^ had reduced relative effect sizes (**Supplementary Fig. 5E**, Welch’s t-test P=1.7e-4).

Negative control sgRNAs are necessary to calibrate the null phenotype and test significance. Current screens employ either non-targeting sgRNAs, which do not target anywhere in the genome, or safe-targeting sgRNAs^44^, which target regions presumed to be biochemically inactive. Previous growth screens with Cas9 nuclease suggest that safe-targeting sgRNAs have stronger effects than non-targeting sgRNAs due to the effects of DNA damage^44^. In contrast, we did not observe a significant difference in the average effect of non-targeting versus safe-targeting sgRNAs in a *CD164* CRISPRi HCR-FlowFISH screen that included 1,000 of both types of negative controls (Welch’s t-test, P value=0.23). However, safe-targeting sgRNAs had significantly greater variance, demonstrating that they are more stringent controls for significance testing (**Supplementary Fig. 6A**, variance of safe-targeting sgRNAs=1.17 or non-targeting sgRNAs=0.86, Levene’s test P value<0.001). While a greater number of negative control sgRNAs reduces their variance, there is no statistically significant difference in the variance of 700 safe-targeting controls compared to all 1,000 (**Supplementary Fig. 6B**). Thus, we recommend including at least this number of safe-targeting sgRNAs, and encourage the use of a common set of safe-targeting sgRNAs (**Supplementary Tables 8, 9**) to allow direct, robust comparisons across screens^44^. These safe-targeting sgRNAs were designed based on Roadmap Epigenomic data, and may intersect activation-associated epigenetic features in a novel cell type or sample–thus, we advise ensuring that the selected safe-targeting sgRNAs target epigenetically inert regions.

Finally, sufficient numbers of sgRNAs targeting the measured gene’s promoter, or the promoter(s) of genes known to regulate the measured phenotype, should be included as positive controls to ensure that strong perturbations can be sensitively detected, and to establish the upper bound of measurable effect sizes. These can be selected from established, genome-wide, gene-targeting CRISPRi libraries^44–46^. We compared the average effects of the 10 sgRNAs closest to each FANTOM and refGene TSSs for the HCR-FlowFISH genes, along with the 4-10 sgRNAs from the human CRISPRi Dolcetto^46^ or hCRISPRi-v2^45^ libraries that were also included in our libraries–these sgRNAs often target one or more TSSs for a given gene and are selected for features that improve on-target efficacy. We found that sgRNAs from the Dolcetto or hCRISPRi-v2 libraries provided average effects similar to the maximum average effect from perturbing all of the FANTOM and/or refGene TSS(s) for 12 out of 14 genes (**Supplementary Fig. 6C**). However, for the gene *FADS2*, there were >2-fold greater effects at some FANTOM and refGene TSS(s) than with the published sgRNAs. Given that neither Dolcetto nor hCRISPRi-v2 was consistently best, including sgRNAs from both published libraries increases the likelihood of having potent positive controls, but designing 10 sgRNAs nearest every TSS–where space allows–maximizes it.

To facilitate design of sgRNA libraries in accordance with these recommendations, we provide a summary of common sgRNA design tools and highlight the specific requirements, advantages and disadvantages, and experimental parameters for each (**Supplementary Table 10**). As a resource, we used GuideScan2^47^ to design sets of sgRNAs with and without filters (GuideScan2-aggregated CFD specificity score ≥0.2 and no ‘TTTT’ sequences) for all human and mouse ENCODE SCREEN^6^ cCREs (**Supplementary Fig. 7, Supplementary Table 8, Supplementary Section 3**). These sets include at least 10 sgRNAs for targeting 85% and 60% (without and with filters, respectively) of the 249,464 human proximal enhancer-like cCREs and 86% and 70% of the 111,218 in mouse (**Supplementary Table 11**). Importantly, the design guidelines provided here are based on modeling of data produced from experiments that were conducted at similar and sufficient coverage and power. These experimental guidelines are discussed below, and deviations from the following recommendations may require that additional controls or sgRNAs be included per target element.

### Cell coverage and sequencing depth impact CRE detection accuracy and sgRNA dropout

We next interrogated how varying the number of cells per sgRNA (cell coverage) impacts accuracy of CRE identification, using CRISPRi HCR-FlowFISH experiments at the *GATA1* locus (**Methods, Supplementary Table 12**). Three CREs were previously identified and verified in all our CRISPRi screens, including growth screens^10^ (**Fig. 1D, 3D**). We tested whether *positive sgRNA targets* (those contained within the three CREs, N=288) can be distinguished from *negative sgRNA targets* (those outside the three CREs, N=13,444) by their log_2_FC effect sizes, where the coordinates of the three CREs were obtained using the CASA peak caller^9^. At low cell coverage (20x), effect sizes of both sets of sgRNA targets had high variance, leading to limited statistical power for distinguishing positive signals from negative control background (**Fig. 3A**). With increasing cell coverage, the variance of negative sgRNA targets approaches zero (Levene’s test P<2.2e-16 for all pairs of 20 vs. 50x, 50 vs. 100x, and 100 vs. 200x), while the variance of positive sgRNA targets stabilizes for coverages ≥50x (Levene’s test, P value=0.010, 0.87, 0.22 for 20 vs. 50x, 50 vs. 100x, and 100 vs. 200x comparisons, respectively). These variance reductions led to significantly higher precision and sensitivity for distinguishing *positive sgRNA* targets from *negative sgRNA targets*, by their effect sizes, relative to 20x cell coverage (**Fig. 3B**). Further, CASA peak calling with 50x-200x cell coverage resulted in accurate identification of the known *GATA1* CREs, while the 20x data resulted in spurious CRE calls lacking CRE-associated epigenetic marks (**Fig. 3C**). Lastly, with cell coverage of 20x, we observe a high dropout rate (percent of sgRNAs with <10 mapped reads in low- or high-expression sorting bins) of ∼12%, which decreases to less than 1% with cell coverage greater than 50x (**Supplementary Fig. 8**). Based on these strong to moderate *GATA1* CREs, experimental cell coverage of at least 100x should be considered the minimum, although higher coverage is advised when feasible. For example, coverage as high as 11,000x has been used in non-coding growth-based screens^17^.

**Fig. 3.**
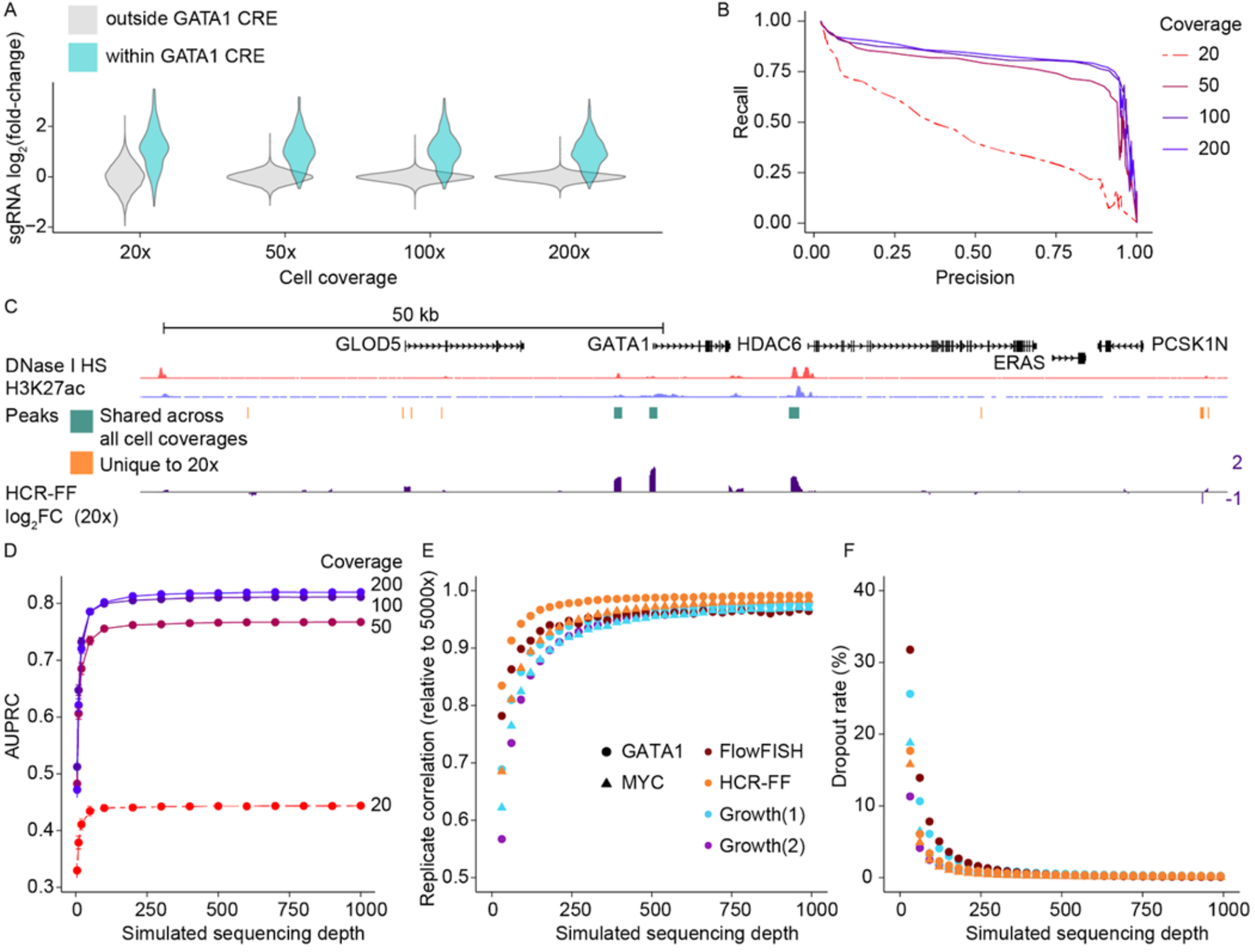
Cell coverage and sequencing depth impact reliable detection of CREs. **A**) Distributions of HCR-FlowFish guidewise log_2_FC effect sizes (total 13,732 PAMs targeted) at various cell coverages, separately for sgRNA targets within (N=288) and outside known *GATA1* CREs (N=13,444). **B**) Precision-recall curve for identifying *GATA1* CRE-targeting sgRNAs using effect sizes from various cell coverages (AUPRC: 20x=0.44, 50x=0.77, 100x=0.81, 200x=0.82; CRISPRi HCR-FlowFish). **C**) log_2_FC signals for 20x and CASA peak calls shared across all coverages and unique to 20x **D**) AUPRC for identifying *GATA1* CRE-targeting sgRNAs with varying sequencing depth (bootstrap sampled) and cell coverages (20x, 50x, 100x, 200x). Dots and error bars indicate averages and 99% confidence intervals over 10 bootstrap samples. **E**) Biological replicate reproducibility (guidewise log_2_FC Pearson correlation), normalized to 5000x sequencing depth and **F**) guide dropout rate (dropout defined as <10 mapped reads) in diverse CRISPRi screens with varying sequencing depth (bootstrap sampled). Dots show an average of over 100 bootstrap samples. Growth datasets are (1) Tycko et. al. 2019 and (2) Fulco et. al. 2019.

We also sought to derive sequencing depth guidelines for non-coding CRISPR screens. We bootstrap sampled on average 5x to 1000x sequencing reads per sgRNA across each of the 20x, 50x, 100x, and 200x cell coverage *GATA1*-locus HCR-FlowFISH screens and assessed normalization strategies (**Fig. 3D, Supplementary Fig. 9, Methods**). With 250x sequencing depth or higher, accuracy (AUPRC for distinguishing *positive* and *negative sgRNA* targets by log_2_FC effect sizes) of HCR-FlowFISH screens for *GATA1* CREs is limited by cell coverage, such that further increase in sequencing depth only marginally improves accuracy. We repeated the analysis in five other CRISPR screens, including growth screens performed at *GATA1* and *MYC* loci, finding 250x sequencing depth was a reasonable minimum for CRE identification accuracy. Further, we observed saturations of biological replicate correlation of guide effects and of guide dropout rate starting at 250x sequencing depth (normalized bootstrap avg. bio-replicate log2FC R>0.9 & average dropout rate <2%, for all screens, **Fig. 3E,F, Supplementary Fig. 10**). Overall, our analyses suggest a sequencing depth of at least 250x for similar CRISPRi screens.

### CASA provides more conservative CRE calls than other analysis methods

Non-coding CRISPR screens can produce noisy results when individual targeting sgRNAs generate variable effects in a genomic interval (**Fig. 4A**). Multiple analysis approaches, or ‘peak callers’, aggregate individual sgRNA measurements from dense tiling screens to nominate CREs. We investigated the use of five peak callers: element-level aggregation of DESeq2 (aggrDESeq2), CASA, CRISPR-SURF, MAGeCK, and RELICS^9,48–51^. The underlying statistical models, input parameters, ease of use, and output formats vary with each method (**Methods, Supplementary Table 13**). We benchmarked the identification of *GATA1* CREs using a CRISPRi tiling growth screen, excluding confounding low-specificity sgRNAs (GuideScan aggregated CFD > 0.2) (**Fig. 4**). While a comprehensive, fully validated, ground truth CRE set is lacking, these CREs have been rigorously epigenetically profiled and studied across multiple functional characterization assays^9,10,12,15^.

**Fig. 4.**
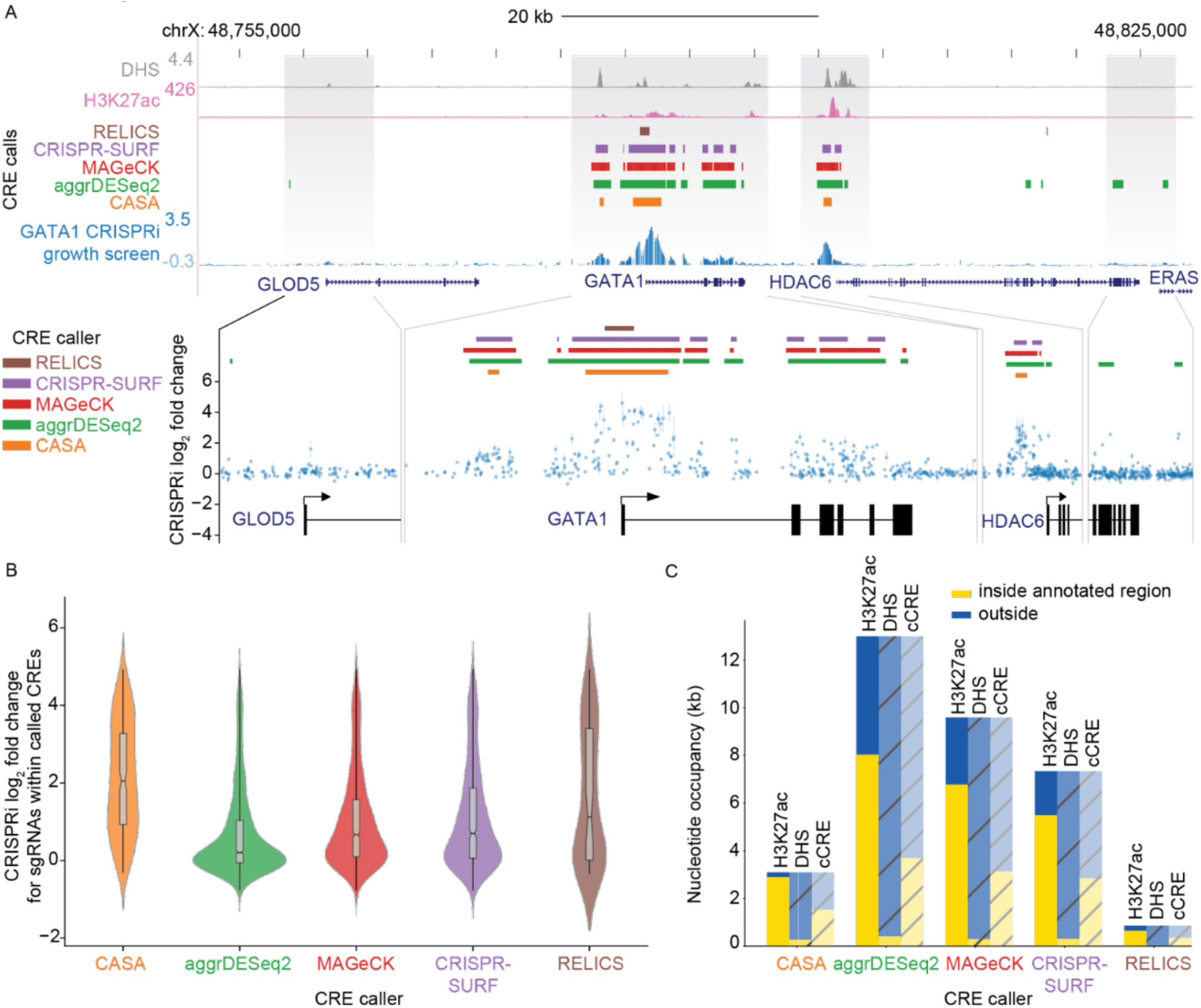
CRISPR screen analysis tools identify CREs with varying selectivity. **A**) sgRNA mediated growth effects (blue), H3K27ac-ChIP signal (pink), and DNase Hypersensitivity signal (gray) for a CRISPRi growth screen at the *GATA1* locus. sgRNAs are filtered to remove any low-specificity sgRNAs (GuideScan aggregated CFD<0.2) which could cause confounding off-target toxicities. Dense tracks show peak calls using 5 different CRISPR screen analysis tools: CASA (orange), aggrDESeq2 (green), MAGeCK (red), CRISPR-SURF (purple), and RELICS (brown). Zoomed- in regions show log_2_FC of individual sgRNA effects (points, mean; bars, min-max range of observations between n=2 replicates). **B**) Distribution of average guide effects calculated from two experimental replicates for sgRNAs falling within peaks identified by different CRISPR screen analysis tools (center line, median; notch confidence interval of the median; box limits, first and third quartiles; whiskers range of all data points; violin, kernel density estimation). **C**) Total peak area inside and outside of annotated chromatin features and ENCODE SCREEN cCREs for each peak caller.

All peak callers nominate the promoter for *GATA1* (**Fig. 4A**) as an active element. Additionally, CREs called by all five peak calling methods corresponded with significantly higher guide effects than shuffled control elements (**Fig. 4B**, P≤2.2e-16, Welch’s t-test), and CREs supported by at least four callers are also supported by chromatin accessibility or H3K27ac signal (**Fig. 4A**). However, the total number of CREs varied across each method with aggrDESeq2 identifying the greatest number, and CASA and RELICS identifying the least number of CREs (aggrDESeq2: n=21, CASA: n=3, CRISPR-SURF: n=14, MAGeCK: n=10, RELICS: n=3). To assess the accuracy of each approach, we quantified the number of nucleotides within the proposed CREs that overlap annotated ENCODE SCREEN cCREs, H3K27ac peaks, and DHSs (**Fig. 4C**). CASA, CRISPR-SURF, and MAGeCK, identified CREs with a greater proportion of nucleotides overlapping these annotations while aggrDESeq2 CREs yielded the largest total overlap but also identified a greater proportion of CREs outside of these annotations. We then measured pairwise Jaccard Similarity between the set of nucleotides nominated by each of the five peak callers and three previously identified *GATA1* regulatory elements (**Supplementary Fig. 11A**). We found that these canonical elements are most similar to CASA and RELICS CREs and least similar to aggrDESeq2 CREs. Finally, we inspected the intersection of *GATA1* CRE calls from each method and found CASA was the only peak calling method that lacked unique, and potentially spurious, *GATA1* CRE calls (**Supplementary Fig. 11B**).

To determine the susceptibility of each peak caller to potential sgRNA off-target effects, we re-analyzed the *GATA1* screen with low-specificity sgRNAs included (**Methods; Supplementary Fig. 12A-D**). Without filtering, the total number of CREs called by aggrDESeq2 increased by >3x (21 CREs vs 68 CREs), and that method identified the greatest number of CREs not identified by other methods, suggesting this approach is particularly sensitive to false positives caused by off-target effects. The total number of CREs called by CRISPR-SURF, MAGeCK, and RELICS increased by 12, 4, and 2, respectively. In contrast, the number of CREs identified by CASA did not change. Taken together, these results support CASA as the preferred method for CRE calling as it may be less affected by off-target effects of low specificity sgRNAs and its CRE calls exhibited the greatest proportion of overlap with CRE-associated annotations. To facilitate future analytical development and benchmarking, we propose processed data file formats that capture critical experimental parameters and include sgRNA-level and CRE-level effect quantification (**Supplementary Sections 4, 5**).

### Perturbation dynamics affect screen sensitivity

Our integrated dataset provides an opportunity to investigate possible interactions between perturbation timing, sgRNA effect sizes, and phenotyping strategy, which remain largely uncharacterized and variable across non-coding screens. Conceptually, a high effect-size sgRNA would be expected to display detectable phenotypic impacts sooner than a weaker-effect size sgRNA, rendering measurement timing a critical aspect of experimental design.

To explore how CRISPR perturbation dynamics and experimental design affect measured phenotypes, we analyzed data from several time points from a CRISPRi growth screen at the *GATA1* locus, in which we delivered sgRNAs by lentivirus into a cell line constitutively expressing the CRISPRi machinery. There is no clear consensus on if the initial plasmid pool of sgRNAs or an early time point post lentiviral-delivery should be used as the comparator to an endpoint sample when computing sgRNA effects. We sequenced sgRNAs in the pre-delivery plasmid pool (plasmid), at seven days after lentiviral guide delivery to cells (T7, the earliest time after selection for sgRNA delivery), and at an endpoint after twenty-one days (T21), (**Fig. 5A**). Comparing plasmid to T7, we observe a significant CRE at the promoter, but do not identify the distal eGATA1 and eHDAC6 CREs (**Fig. 5B**). However, both distal CREs are identified in the plasmid-T21, or T7-T21 comparisons (**Fig. 5B**) and the peak at the promoter widens by ∼1 kb with increasing sgRNA effect sizes.

**Fig. 5.**
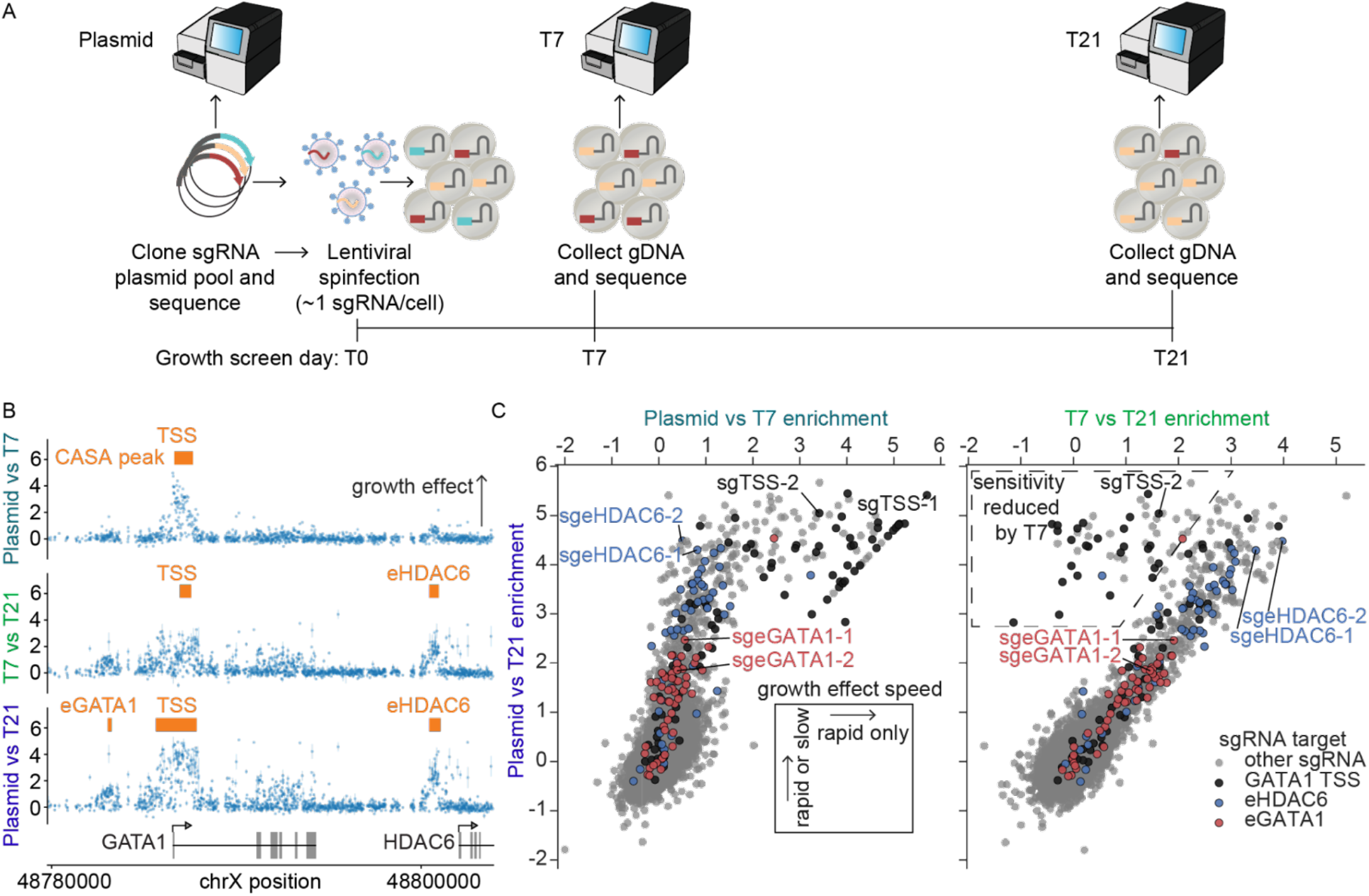
Perturbation dynamics impact screen sensitivity and resolution. **A**) Timeline of CRISPRi growth screen with quantified sgRNA abundances of the sgRNA plasmid library pre-delivery, and 7 and 21 days after sgRNA lentiviral delivery. **B**) CRISPRi growth screen at *GATA1* locus shown with varied time point comparisons (top: Plasmid vs T7, middle: T7 vs T21, bottom: Plasmid vs T21) used to compute sgRNA effect sizes. Each dot shows the average log_2_(fold-change) effect size of two biological replicates for an sgRNA, and the error bar shows the range. CASA peak calls for significant growth effects are shown. The *GATA1*-regulating CREs eGATA1, *GATA1* TSS, and eHDAC6 are labeled with their corresponding CASA peak calls. **C**) Scatter plot of sgRNA effect sizes as determined by varied time point comparisons. Each dot shows the average of two biological replicates for an sgRNA. Black or colored dots are sgRNAs targeting the TSS or enhancers, respectively. The sgRNAs along the diagonal line of points, including sgTSS-1, drop out by T7 and thus are absent from the T7 vs T21 comparison. sgRNAs selected for validation assays are labeled (**Supplementary Table 14**).

While the sgRNA effect sizes from these two time point comparisons are correlated (R^2^=0.71), a subset of sgRNAs (<1%) displayed time point-dependent effects (**Fig. 5C**). These sgRNAs are strong (log_2_FC>3) in a plasmid-T21 comparison, but have reduced effect sizes in a T7-T21 comparison (see dashed line bounded region in **Fig. 5C**, right panel). These sgRNAs largely target the *GATA1* TSS. One of these sgRNAs (sgTSS-2) has been individually validated to reduce *GATA1* expression and growth (**Supplementary Fig. 2C and Supplementary Table 12**). Another validated sgRNA (sgTSS-1, **Supplementary Fig. 2C**) displays the third strongest effect (out of 9,923 sgRNAs) in the plasmid-to-T21 comparison (log_2_FC=5.4) and the strongest effect in the plasmid-to-T7 comparison (log_2_FC=5.7), but this sgRNA drops out by T7 and is not observed in the T7-to-T21 comparison thus becoming a false negative possibly due to the rapid nature of its effect. Together, this suggests these rapidly depleted sgRNAs can cause bonafide growth phenotypes, and the strongest hits may be most affected by reduced sensitivity in the T7-to-T21 comparison.

We reasoned that screens based on complex cell phenotypes, such as growth, may be more sensitive to perturbation dynamics than screens that directly readout transcriptional changes, given the need for sufficient time for gene expression to manifest impact on the downstream phenotype. Indeed, an HCR-FlowFISH screen of *GATA1*, in which sgRNA abundances were compared pre- and 2 days post-CRISPRi induction by doxycycline, identified both the promoter and the two distal CREs (**Fig. 1D**). This screen format was not susceptible to reduced power to detect the strongest TSS-targeting sgRNAs, as was observed when comparing T21 and T7 in the growth-based readout. Together, we suggest comparisons to plasmid sgRNA abundance before starting phenotypic selection, for example by measuring sgRNA abundance in the input plasmid library or in cells before CRISPRi expression in an inducible system.

### CRISPRi effects in the gene body are strand-specific

Most CRISPR screens model and analyze sgRNA effects without considering the potential impact of which DNA strand is targeted. However, strand independence of CRISPRi perturbations has not been empirically assessed. Analyzing a CRISPRi growth screen tiling *GATA1*, we surprisingly found sgRNAs targeting the coding strand affected growth while template-targeting sgRNAs did not (P-value<1e-15, **Fig. 6A**). This difference was only observed in the gene body, perhaps related to RNA Pol II binding the template strand during gene transcription. We again observed significantly greater effects for sgRNAs targeting the coding strand within the gene body in the *FADS1* and *FADS2* HCR-FlowFISH CRISPRi tiling screens (P value<1e-15, **Fig. 6B**). These coding-strand effects were uniform throughout the transcribed gene body and ended at the transcription end site (**Supplementary Fig. 13A**). Notably, strand-bias was not observed for sgRNAs targeting the promoters (**Fig. 6A-C**). We did not observe such effects from the same library of sgRNAs targeting either strand in the gene body when using dCas9 alone (**Fig. 6A**), or when using CRISPRa (**Fig. 6D**), suggesting this phenomenon depends on the KRAB repressor and not solely on dCas9 binding (**Fig. 6.E**)

**Fig. 6.**
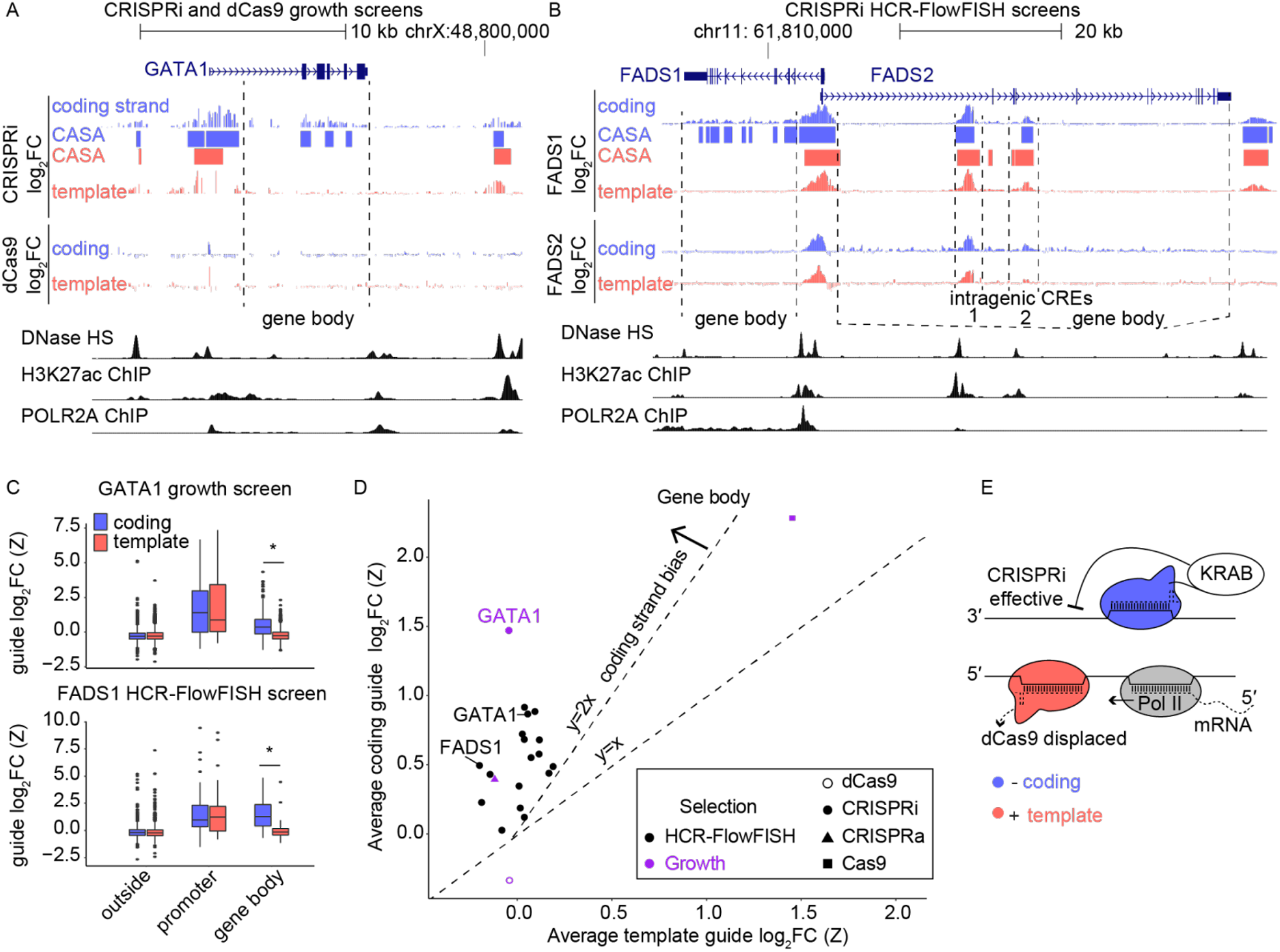
CRISPRi effects in the gene body are strand-specific. **A**) Strand-specific CRISPRi growth screen effects tiling *GATA1*. CRISPRi and dCas9 tracks show the average of two biological replicates, comparing Day 21 to plasmid (N=2,541 coding- and 2,263 template-strand targeting sgRNAs). **B**) Strand-specific CRISPRi HCR-FlowFish screen effects tiling FADS1 and FADS2. CRISPRi tracks show the average of two biological replicates, comparing high-versus low-expression bins for the target gene (N=4,609 and 4,942 sgRNAs per strand). **C**) Distributions of sgRNA effects (average of 2 screen replicates) in the gene body, and at the promoter (within 2 kb upstream of TSS), when sgRNAs are categorized by target strand, in the (top) *GATA1* CRISPRi growth screen (n=2026, 1731, 34, 27, 100, and 77 sgRNAs from left to right boxes) and the (bottom) FADS1 HCR-FlowFish screen (n=3121, 3249, 90, 69, 520, and 702 sgRNAs). Boxes show the quartiles, with a line at the median, vertical lines extend to 1.5 times the interquartile range, and dots show outliers. Asterisk denotes significance with P<1e-15 by T-test. **D**) Strand-specificity across screens tiling 17 loci for sgRNAs targeting the gene body. Each point is the average effect of all sgRNAs from a screen targeting that region, averaged across two screen biological replicates, with color indicating the phenotypic readout, and shape indicating the type of CRISPR perturbation. **E**) Proposed model of gene body strand bias, wherein dCas9 binding could be reduced on the template strand due to competition with Pol-II-mediated transcription, rendering KRAB ineffective. In contrast, when targeting the coding strand, KRAB can be effective.

To determine if this effect was present more generally, we expanded our comparison to 17 additional experiments (**Methods**). In all 17 CRISPRi screens, the average effect sizes of sgRNAs targeting coding strands within gene bodies were more than 2-fold higher than those targeting the template strands (**Fig. 6D**). In contrast, for all 17 corresponding promoters, there was no difference between coding and template strands (**Supplementary Fig. 13B**).

Many enhancers reside within gene bodies^52^, motivating us to consider if these CRISPRi effects throughout gene bodies could be distinguished from effects at intragenic enhancers. *FADS2* contains intragenic enhancers, as determined by concordant signals from CRISPRi HCR-FlowFISH, DHS, and H3K27ac ChIP-seq (**Fig. 6B**). In contrast to elsewhere in the gene body, sgRNAs targeting both strands in these two enhancers had a significant effect on FADS2 expression, although sgRNAs targeting the coding strand had a moderately greater effect than those targeting the template strand (P value=0.034 and 0.018 respectively, **Fig. 6B** and **Supplementary Fig. 13C**). Practically, these results demonstrate the necessity of considering strand to reliably identify intragenic CREs with CRISPRi: in CREs there are strong effects from sgRNAs targeting both strands, throughout the rest of the gene body there are subtle effects from coding strand-targeting sgRNAs only.

## Discussion

Multiple CRISPR-based methods have been employed to examine cis-regulatory mechanisms yet there has not been a consensus on experimental design parameters and analysis methods. Differences in the design, execution, and analysis of CRISPR screens and lack of common controls render a systematic evaluation of methodologies difficult, especially comparisons of screen sensitivity and specificity. To address these limitations, we performed a comprehensive analysis of the ENCODE non-coding CRISPR screen datasets and proposed guidelines for successful assay implementation, standardized file formats, and processed data expectations. Implementing the guidelines proposed here will facilitate further cross-screen comparisons, meta-analyses, and improved power for CRISPR screens.

The ENCODE4 Functional Characterization Centers identified 887 CREs, but these experiments were performed in a limited set of biosamples and represent a small fraction of the true number of functional CREs. The identified CREs may be biased by the library designs, phenotypic readouts, specific genomic loci perturbed, and analysis methods used in these experiments. More systematic screening with high-quality design, execution, and analysis across multiple phenotypes and genomic regions is needed to capture the full range of cis-regulatory mechanisms. Building a larger, diverse collection of “gold-standard” CREs will improve guidelines for selecting sgRNAs at elements overlapping non-accessible but otherwise active (e.g. by H3K27ac) regions, and will empower refinement and benchmarking of methodological guidelines and analysis techniques, especially for characterizing CREs of weak effect.

cCRE-targeting screens can interrogate thousands of perturbations at once, but lack full-genome saturation scale. Balancing sensitivity and specificity, especially for modest effect size CREs, with screening scale requires rational, evidenced based sgRNA library design. One important future design consideration is our observation that the strongest sgRNAs are nearest to distal CRE DHS summits. This can be potentially attributed to increased accessibility improving CRISPRi efficiency and/or higher transcription factor motif density, but we note additional modeling will help determine an sgRNA sequence’s contribution to on-target efficacy.

We observed a previously unreported CRISPRi strand bias specific to gene bodies, which correlates with previous findings that gene body transcription displaces Cas9 nuclease^53^. Whereas others have recommended use of template strand-targeting sgRNAs with Cas9, reasoning that Cas9 displacement could be productive for genome editing, our results show coding strand-targeting guides are stronger for epigenome editing with CRISPRi. We suggest using strand-aware analysis to distinguish intragenic CREs from the subtle effects of CRISPRi throughout the gene body. Future research could illuminate how CRISPRi in the gene body, but outside of CREs, mechanistically impacts gene expression.

We compared several peak callers for *de novo* CRE discovery in tiling screens and found that while they all identify positive control CREs, CASA maintained both sensitivity and precision while others were more prone to false positives from off-target noise. In both sparse, cCRE-targeting and cCRE/locus-tiling screens, including biological replicates and increasing sgRNA number were critical for detecting weak elements and improving power. We note that even for strong CREs, many more sgRNAs are needed than often used in current screens (suggested minimum: 10 sgRNAs within 100 bp of the DHS peak). We advise considering the guidelines described in this study (i.e. experimental coverage, peak caller comparisons, sgRNA numbers per element) as minimums, and empirically evaluating power in new experimental systems. We expect future analytical packages could incorporate replication, strand-bias, and sgRNA efficacy to improve CRE detection.

Identifying functional CREs with perturbation experiments is an imperative step towards understanding the mechanisms that govern gene regulation and how disruption of these CREs contribute to disease. However, optimal experimental and analytical parameters are needed to increase the scale and/or sensitivity of CRISPR screens, especially as they are increasingly applied with multiplexed readouts and in single-cell schemas. By providing specific, evidenced-based recommendations based on a diverse set of CRISPR screens in the ENCODE database, along with suggested sgRNAs for cCREs, this work will accelerate the functional characterization of regulatory elements genome-wide and make non-coding CRISPR screening methods accessible to the broader community.

## Notes

### Competing Interest Statement

A.K. is scientific co-founder of Ravel Biotechnology, is on the scientific advisory board of PatchBio, SerImmune, AINovo, TensorBio and OpenTargets, is a consultant with Illumina and owns shares in DeepGenomics, Immuni and Freenome. C.A.G. is a co-founder of Tune Therapeutics and Locus Biosciences, and an advisor to Tune Therapeutics and Sarepta Therapeutics. C.A.G. is an inventor on patents and patent applications related to CRISPR epigenome editing. J.T. and M.C.B. acknowledge an outside interest in Stylus Medicine. L.L. is currently employed by Sana Biotechnology. P.C.S is a co-founder of and consultant to Sherlock Biosciences and Board Member of Danaher Corporation. She is a shareholder in both companies. W.J.G. is a co-founder of Epinomics and an adviser to 10X Genomics, Guardant Health and Centrillion.

