## Supplementary Information, Figures, and Methods. for "Multi-center integrated analysis of non-coding CRISPR screens"

Supplementary Material Related to:
***Multi-center integrated analysis of non-coding CRISPR screens***

### Table of contents

**Table of contents** 1
**Supplementary Sections** 3
Supplementary Section 1. Navigating CRISPR Screening Data on the ENCODE Portal 3
Supplementary Section 2. Validation of CREs identified from CRISPR screens 6
Supplementary Section 3. Design of sgRNA libraries targeting all ENCODE SCREEN cCREs 7
Supplementary Section 4. Standardized non-coding CRISPR Screen File Formats 8
Supplementary Section 5. Processed non-coding CRISPR screen file formats 9
Supplementary Section 6. sgRNA Sequence and Coordinate Mapping 10

**Supplementary Figures** 11
Supplementary Fig. 1. Meta-analysis of K562 screens nominates features of functional CREs. 11
Supplementary Fig. 2. Individual sgRNA validations correlate with non-coding CRISPR screen results 12
Supplementary Fig. 3. Analysis of CRISPR screens at the MYC and GATA1 loci. 13
Supplementary Fig. 4. Selecting cCREs and targeting sgRNAs near DHS summits 14
Supplementary Fig. 5. Analysis of sgRNA specificity and power related to sgRNAs per element 15
Supplementary Fig. 6. Evaluating methods of selecting negative and positive control sgRNAs 16
Supplementary Fig. 7. GuideScan2-designed sgRNAs targeting all cCREs from the ENCODE SCREEN portal. 17
Supplementary Fig. 8: Guide drop out rate with varying HCR-FlowFISH sorting depths 18
Supplementary Fig. 9: Comparison of effect size normalization methods 19
Supplementary Fig. 10: Representative bootstrap samples for low and high sequencing depths 20
Supplementary Fig. 11. Overlap between peak calls on specificity-filtered CRISPRi tiling screen of GATA1 locus 21
Supplementary Fig. 12. Peak calls without filtering out low specificity sgRNAs 22
Supplementary Fig. 13. CRISPRi strand bias in the gene body 23
Supplementary Fig. 14. Mapping sgRNAs to reference genome for data standardization 24
Supplementary Fig. 15. ‘Functional Characterization’ card on the ENCODE home page 25
Supplementary Fig. 16. Filtering search results in the ENCODE portal 26
Supplementary Fig. 17. An experiment series summary page 27
Supplementary Fig. 18. The ‘Files Section’ of the experiment series summary page 28

**Supplementary Tables** 29

**Materials and Methods** 30
Cell lines and cell culture 30
1.1 The ENCODE CRISPR Screen Database and overlap with cCREs 30
1.2 CRISPR screen comparisons with individual sgRNA validations 31
1.3 Cross-screen analysis at GATA1 and MYC 31

2.1 Evaluating sgRNA effects in DHS or H3K27ac peaks 31
2.2 Evaluating sgRNA effects as a function of distance from the DHS summit 31
2.3 Effect size-dependent sgRNA number per element power analysis 32
2.4 Off-target sgRNA enrichment analysis 32
2.5 Safe vs. non-targeting negative control variance statistical analysis 32
2.6 Promoter-targeting “positive control” sgRNA selection analysis 32
3.1 Cell coverage / sorting depth titration experiments for HCR-FlowFISH 33
3.2 Bootstrap sampling analysis for simulating CRISPR screens performed at various sequencing depths 33
4. Peak caller comparisons 34
5. Comparison of timepoints 35
6. Strand specific quantification of sgRNA effect sizes 35
Data availability 36
Code availability 36
Public datasets accessed 36
Author contributions 36
Acknowledgements 36
Conflict of Interest Statements 37

### Supplementary Sections

**Supplementary Section 1. Navigating CRISPR Screening Data on the ENCODE Portal**

The ENCODE portal has been updated to support submission and subsequent navigation of functional characterization assay data allowing users to navigate these data along various facets: biosample, organism, readout and target, perturbation modality and scale of perturbations, and genome build. In total, the portal contains 119 CRISPR screen experiments, comprising 106 experiments performed in human cell lines and 13 experiments performed in mice as of August 2022 (**Supplementary Table 1**). The most recent data release (November 2022) increased the total number of experiments to 129 (116 in human cell lines, 13 in mice) and we anticipate the release of additional experiments by the end of 2022.

The navigation guidelines covered here allow rapid exploration and filtering of functional characterization datasets generated and made available by the ENCODE Consortium. It guides users through the ENCODE portal to search, visualize and download experiment series data using a web browser (**Supplementary Fig. 15-18**). The functional characterization datasets available in the portal were generated by assays which study the relationship between a DNA sequence and its regulatory activity, such as Massively Parallel Reporter Assays and CRISPR screens.

Find Functional Characterization datasets on the portal

1. Navigate to the ENCODE portal home page at https://www.encodeproject.org/.

2. Locate and click on the “Functional Characterization" card (**Supplementary Fig. 15**).

3. The search result page lists the available functional characterization data. As of August 2022, the ENCODE portal had 477 functional characterization datasets listed (note that only 25 are listed on the page by default)

4. Filter the search results :

    - The sidebar on the left side of the search results page is populated with facets that allow users to filter search results using different criteria. Locate the “Quality” facet group and click on it, notice that the “released” entry under the “Status” facet is highlighted in blue, indicating that the search results have been filtered for datasets that have been released to the public.

    - Selections can be applied to several facets at a time and the combined filters possess an “AND” relationship. Scroll back up to the facet group “Provenance” and under the facet “Lab” select “Pardis Sabeti, Broad”. Now the search results have been filtered to include only functional characterization datasets that have been performed in Dr. Sabeti’s laboratory AND have been released to the public.

    - Review the list of search results. As of August 2022, the facets selections above returned 20 series datasets. Select the dataset ENCSR408VHJ by clicking on its title “CRISPRi Flow-FISH screen in K562 with HCR-FlowFISH readout of MYB” (**Supplementary Fig. 16**).

Navigate Experiment Series Summary Page

The experiment series summary page is organized in six distinct sections: (A) Page Title, (B) Summary, (C) Attribution, (D) Experiments, (E) Control Experiments and (F) Files section (**Supplementary Fig. 17**).

- The “Summary” section (**Supplementary Fig. 17B**) contains key information about the series, including but not limited to donor, assay, biosample summary, diseases, treatments. It will also include a link to the elements reference dataset that contain information about the investigated functional elements or genomic loci.

- The “Attribution” section (**Supplementary Fig. 17C**) lists the lab, award and project. It will specify if there are aliases used for the series and if there are cross-references to the dataset in other public repositories (external resources).

- The “Experiments” section (**Supplementary Fig. 17D**) lists in a table all functional characterization experiments that have been included in the series. The first column of the table provides the accessions of the experiments, followed by the assay type of the experiment. For experiments that measure expression readout of a specific genomic locus, the column “Examined loci" will be populated with the relevant information.The next column includes biosample summary, followed by columns listing the lab and status information. Additional information about the various statuses of the experiments and other objects on the portal is available at https://www.encodeproject.org/help/getting-started/status-terms/. The cart option is the last column in the Experiments section table. It allows grouping of individual experiments using the cart mechanism. Note, that you need to create an ENCODE portal user account to take advantage of the cart features. Clicking the accession of a particular functional characterization experiment in the table of the “Experiments section” will take you to the summary page of that experiment. It provides further information on the associated metadata and links to the various experimental components.

- The “Control Experiments” section (**Supplementary Fig. 17E**) lists the accessions of all auxiliary and control experiments that belong to the experiment series. The first column in this section provides the accessions of the control experiments, followed by the control type column. The control experiments have different columns in comparison with the columns of the table in the Experiments section. The descriptive biosample summary is applicable in one (ENCSR692ZUM), but not in the other (ENCSR206MVL) control experiment. ENCSR206MVL is an auxiliary experiment that contains the sequencing result of the cloned sgRNA library. There is no biosample associated with this auxiliary experiment, hence the empty biosample summary. ENCSR692ZUM represents the “base-line” CRISPR screen control experiment and its biosample summary is provided. The lab and status columns provide information for each of the control experiments in the series.

- The “Files” section (**Supplementary Fig. 17F**) is the final section of the experiment series summary page. This section is divided into three tabs: Genome browser, Association graph, and File details.

- The “Genome browser” tab (**Supplementary Fig. 18A**) provides rapid visualization of tracks using the embedded Valis genome browser. All visualizable tracks are shown by default. Visualized on the top is the reference genome assembly (GRCh38) followed by the gene tracks (GENCODE V29) and a track with the SNPs reported in The Single Nucleotide Polymorphism Database (dbSNP). The next two tracks are ENCODE specific tracks showing the latest version of the representative DNase hypersensitivity sites (rDHSs) and candidate cis regulatory elements (cCREs) that are a result of integrative analysis of the ENCODE consortium data. The next track visualizes the guide RNAs locations within the examined region with functional elements specified in the Elements reference dataset (ENCSR827WZZ) associated with the functional experiments in the series. The last two tracks in the genome browser visualize the perturbation signal from the two Flow-FISH CRISPR screens included in this series.

- The “Association graph” tab (**Supplementary Fig. 18B**) displays the data provenance and derivation of the various processed files. The nodes in the graph are clickable and display more information about the node. The yellow nodes represent files, while the blue nodes represent steps in the computational analysis. Click on the yellow node to view the file’s accession and other metadata such as file type, output type, mapping assembly, lab and submission date. Click on the blue node to view information about the relevant pipeline analysis step, the step type, the inputs and outputs, and the software used.

- The “Files details” tab contains several collapsible sections (**Supplementary Fig. 18C**). Each section lists data files that are associated with the dataset. The files in each section are presented in a table with information about the file’s accession and other metadata such as file type, output type, mapping assembly, size and submission date. Each file is presented in a separate row. A small download icon next to each file accession allows users to download a single file at a time. The collapsible section with the ENCAN792OTU identifier contains files originating from the collective analyses of the experiments in the series. The sections with the experiment identifiers (ENCSR476KHP, ENCSR211TEA, ENCSR206MVL and ENCSR692ZUM) lists files from the corresponding datasets included in this series. The “Reference data" section contains files that capture the investigated loci information.

#### Supplementary Section 2. Validation of CREs identified from CRISPR screens

High-throughput screening approaches enable perturbations of thousands of cCREs but can suffer from false positives due to technical limitations including cell number, low sequencing depth, and variability between replicates. As such, it is critical to validate hits with individual perturbations and assess whether the validation supports the screen result.

To this end, CREs identified in ENCODE CRISPR screens have been validated via individual sgRNA perturbations to regulate the phenotype used in the respective screen using RT-qPCR and growth competition assays[1–5]. Similarly, CRE-phenotype connections from external datasets have been confirmed using RT-qPCR in addition to more complex characterization methods, including siRNA perturbation of promoter CREs to rule out secondary effects of perturbations leading to changes in gene expression[6], drug resistance assays[7], and excising the CRE in vivo[8].

At minimum, individual sgRNAs should be delivered to the same cell line used in the screen to confirm the perturbation's effect on the screening phenotype is reproducible. Since screens are often performed in clonally-derived cell lines, effects of individual perturbations may be clone-specific. To confirm the change in phenotype is not clone-specific, we recommend delivering the sgRNA(s) to at least one other clonally-derived cell line expressing the same effector, a polyclonal cell line expressing the effector, or in combination with the effector to an unmodified cell line, and measuring the screening phenotype. For FACS-based readouts, when possible it is also advised to confirm the effect on the phenotype using an alternative characterization method, such as antibody staining for a screen performed with an endogenously tagged gene. Finally, orthogonal editing methods can be used to further confirm the CRE-phenotype connection. For example, a CRE-gene connection identified using CRISPRi could be confirmed by excising the CRE via Cas9-paired sgRNA deletion. Alternatively, if the CRE sequence is sufficiently conserved in other model organisms (e.g. mice), the region can be perturbed in vivo with more complex downstream phenotypic characterization.

**Supplementary Section 3. Design of sgRNA libraries targeting all ENCODE SCREEN cCREs**

To generate sgRNA libraries targeting all human and mouse ENCODE SCREEN v4 cCREs, agnostic of cell type, we first constructed genome-wide GuideScan2 databases for the most recent hg38 and mm10 patches, excluding alternative chromosomes in our analysis. This resulted in two BAM databases containing off-target information, cutting efficiency and specificity scores. The hg38 BAM database is 159 GB in size with 656 million sgRNAs. The mm10 BAM database is 97 GB in size with 116 million sgRNAs.

Next, for each organism and cCRE type, we downloaded the respective cCRE region BED files from the ENCODE SCREEN v4 Registry and used a custom data structure to map the GuideScan2-designed sgRNAs to the cCREs. Specifically, we loaded cCREs into a chromosome-indexed interval tree to enable efficient sgRNA to cCRE mapping and iterated through sgRNAs in the BAM files to find their corresponding cCREs, if any, then emitted the relevant information. This procedure was fast, taking $O(n*\log m)$ time, where $n$ is the number of sgRNAs in the BAM file and $m$ is the number of cCREs; in contrast, other approaches, such as with samtools, took $O(nm)$ time. With parallelization, we could build the database in <2 hours.

Without any filtering based on sgRNA features, there was a median of 26 sgRNAs per cCRE for human proximal enhancer-like signature (pELS) cCREs and fewer for the other cCRE types (**Supplementary Fig. 7 and Supplementary Table 8**). In addition to the unfiltered, genome-wide cCRE sgRNA libraries, we constructed a filtered version of each based on our guidelines that is amenable to either phenotypic (e.g. proliferation) or transcriptional screening readouts (e.g. HCR-FlowFISH). This filtered version removes sgRNAs with a ‘TTTT’ sequence or a GuideScan2-aggregated CFD specificity score<0.2. For each sgRNA, we computed the distance to its nearest cCRE center, as a proxy for the DHS summit. The sgRNA position was considered the position three nucleotides away from the PAM. As the ENCODE SCREEN cCRE regions are defined by the underlying accessibility signal, and given that DHS signal varies across biosamples even at shared peaks, the center of each cCRE was considered a reasonable cell type-agnostic proxy for the DHS summit. We then sorted sgRNAs by their distance from the DHS summit and selected the closest 20 sgRNAs for each cCRE. This list of filtered and sorted sgRNAs can be downloaded as CSVs from: Guidescan2 ENCODE Results. The source code for this pipeline can be found at: https://github.com/schmidt73/encode_pipeline.

**Supplementary Section 4. Standardized non-coding CRISPR Screen File Formats**

As we found important differences in screen datasets, the ENCODE CRISPR working group has established uniform file formats to standardize sgRNA design, reporting, and analysis, for CRISPR screens. Six data standards are proposed for future experiments, each incorporating crucial CRISPR screen-specific information into existing standard file formats. The minimally required columns capture the necessary and sufficient parameters for interpreting and analyzing a screen, and can flexibly compare sgRNAs from one screen to the next even if there are differences in the guide spacer design. Three files are proposed to describe data at the sgRNA level: A) ***guide locations*** - a .bed/bigBed visualizable file that describes PAM coordinates, B) ***guide quantifications*** - a .tsv minimally processed file with sequencing counts for each guide in a single sequenced sample, and C) ***perturbation signal*** - a .bigWig visualizable file of fully processed guide data displaying effect sizes of individual sgRNAs. Another set of files describe the data at the CRE level: A) ***element quantifications*** - a .bed+ file describing CREs measured in the screen with additional information such as effect sizes and contributing sgRNAs, B, C) ***element gene interactions signal*** and ***element gene interactions P value*** - .bigInteract visualizable files linking cCRE to target gene (if known) with effect size or significance (P value) of interaction. Full specifications of the file formats are provided in **Supplementary Section 5** and **Supplementary Tables 13-14**. Adopting these standardized file formats as a field will enable cross-screen comparison, streamline meta-analyses, and improve overall usability of the data, and as more non-coding screen data becomes available, our standards will accelerate the iterative pace of analysis and discovery.

**Supplementary Section 5. Processed non-coding CRISPR screen file formats**

The six standardized data file formats used in this study are detailed below and provide references and quantification at the individual sgRNA-level and CRE-level.

- Guide_location
  - `.bed` of guide coordinates, and corresponding `.bb`, should be bed6
  - Name and score are optional.
  - Includes all regions targeted in an experiment, i.e. include non-significant regions.
  - One per element reference file.
- Guide_quantification
  - `.tsv`, file format specified in **Supplementary Table 13**.
  - This is a minimally processed CRISPR specific datatype meant to collapse fastqs down to the guide-level, enabling cross experiment meta analyses.
  - One per fastq file.
- Perturbation_signal
  - `.bw`, base pair perturbation signal.
  - This is a fully processed file meant for visualization. Use the z-transformed $\log_2$FC for signal values, with 1st bp ('N' in NGG) as location to plot value. On rare occasions with two values at one base pair, average the values.
  - One per replicate per series.
- Element_quantification
  - `.bed`, bed3 with custom additional columns specified in **Supplementary Table 14**.
  - Validated from `.as` file.
  - This is a fully processed file type meant to represent significant CREs.
  - One per series (for replicating peaks) or one per replicate.
- Element_gene_interaction_signal
  - `.bigInteract`, links CRE (from `Element_quantification` file to target gene (if known) with effect size)
  - This is a fully processed file type meant to visualize the `Element_quantification` file.
  - One per series.
- Element_gene_interaction_pvalue
  - `.bigInteract`, links CRE (from `Element_quantification` file to target gene, if known, with P value)
  - This is a fully processed file type meant to visualize the `Element_quantification` file.
  - One per series.

#### Supplementary Section 6. sgRNA Sequence and Coordinate Mapping

As sgRNA design and synthesis strategies are variable, we found direct comparison of guide sequences poorly reflects commonalities between screens. Instead, guide sequences converted into their PAM coordinates in a common genome build (hg38) (**Methods**) resulted in improved overlap (percent overlap with sequence match vs PAM method for *MYC* and *GATA1*). We also compared the effects of lifting over PAM coordinates between genome builds vs. mapping guide sequences with short read aligners to harmonize screening data, and we observed that, for 3 out of 4 guide library constructed in hg19, bowtie had higher rate of unique and exact mapping of sgRNA to hg38 genome build **(Supplementary Fig. 14)**

#### Supplementary Figures

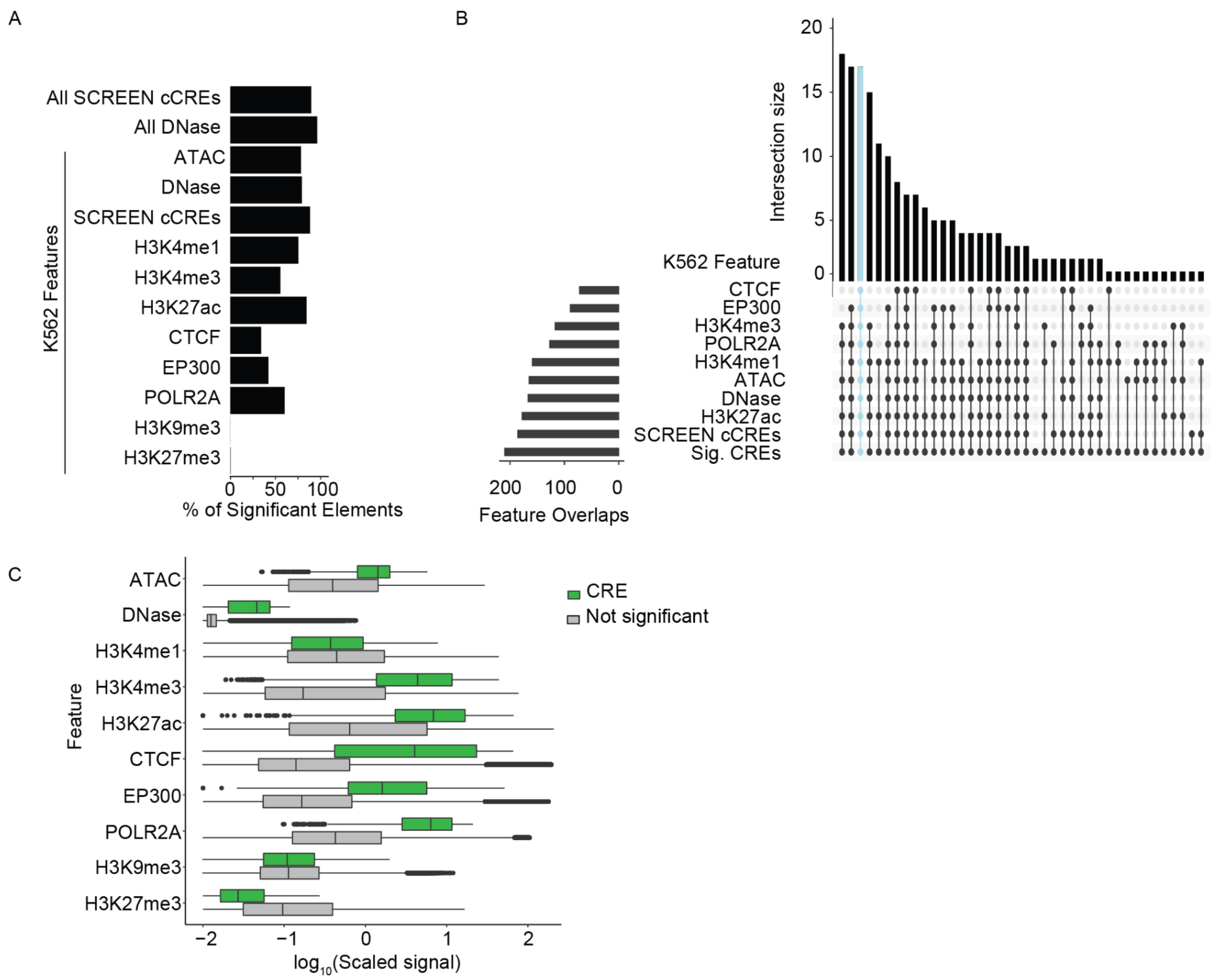


**Supplementary Fig. 1. Integrated analysis of K562 screens nominates features of functional CREs.**

**A**) The percent of total significant CREs (N=210) that intersect union sets of annotations from ENCODE biosamples and K562 annotations. **B**) Upset plot of the intersection of significant CREs with SCREEN K562 cCREs, and K562-annotated accessible chromatin regions, histone marks, EP300, CTCF, and POLR2A, peaks. Blue highlight indicates CREs that intersect all features. **C**) Signal fold change over background for K562 features in CREs (N=210) versus perturbed regions (N=3213). Significant and non-significant perturbation regions colored by green and gray, respectively. Note each value was increased by 0.01 and then $\log_{10}$-transformed for visualization. All comparisons except H3K9me3 were significant at P value<0.01 (Student's t-test). Full test results and mean and median signal values reported in **Supplementary Table 7**. Each box ranges from the first quartile to the third quartile with a line drawn at the median. Lines extend to 1.5x the interquartile range and individual dots extending beyond this range indicate outliers.

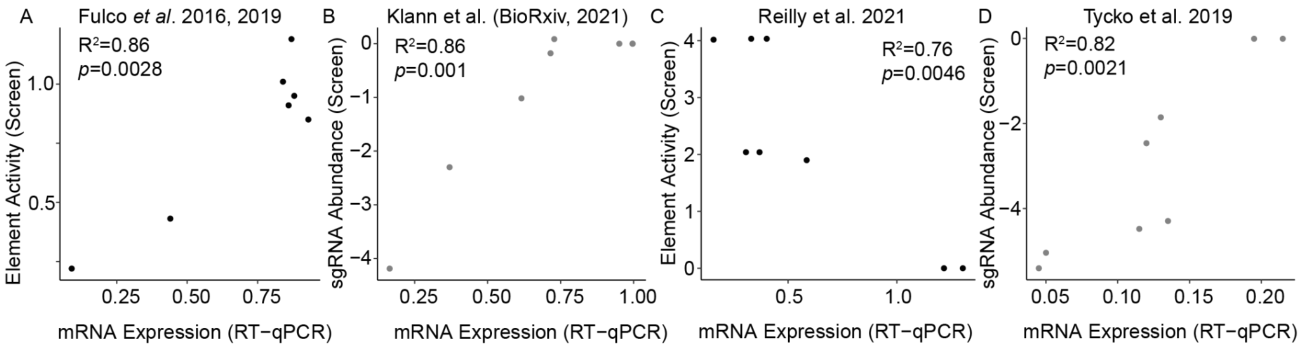


**Supplementary Fig. 2. Individual sgRNA validations correlate with non-coding CRISPR screen results**

**A**) Element-level score from FlowFISH screen in *GATA1* locus versus *GATA1* mRNA expression. A lower element activity value (y-axis) indicates the regulatory element exhibited greater function in the experiment. **B**) Element-level score from FlowFISH screen in FADS1/3 locus versus FADS3 mRNA expression[5]. A greater element activity score indicates the regulatory element exhibited greater function in the experiment. **C**) Abundance of sgRNAs tiling the *GATA1* locus versus *GATA1* mRNA expression[1]. sgRNA labels next to each point correspond to labels in **Fig. 5** and **Supplementary Table 12**. A lower sgRNA abundance indicates the effect of the perturbation was more detrimental to cell growth, and the perturbed element exhibited greater regulatory function. **D**) Validations targeting distal enhancer of LMO2 identified in growth screen targeting all DHSs in K562s[4]. Abundance of sgRNAs versus LMO2 mRNA expression. A lower sgRNA abundance indicates the effect of the perturbation was more detrimental to cell growth, and the perturbed element exhibited greater regulatory function. All validations were performed using lentiviral transduction of individual sgRNAs into dCas9$^{KRAB}$-expressing K562 cells (Pearson correlation values and P values noted within each panel; black and gray indicate FACS-based and growth-based screens, respectively). For **A**,**C**,**D**, each point is the mean of two biological replicates. For **B**, each point is the mean of three biological replicates.

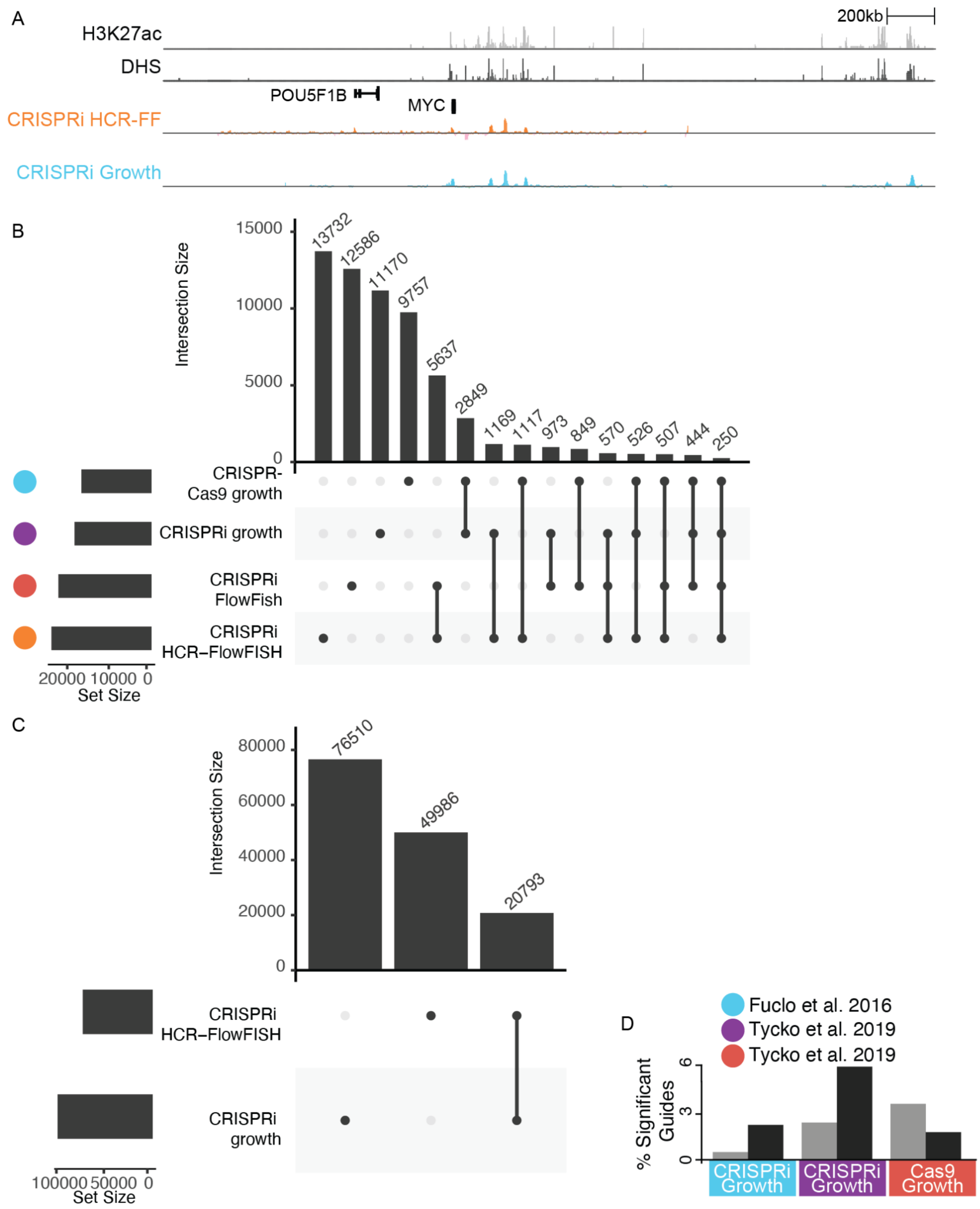


**Supplementary Fig. 3. Analysis of CRISPR screens at the *MYC* and *GATA1* loci.**

**A**) Genome browser snapshot of the *MYC* locus including H3K27ac (light gray) and DHS signal (dark gray) in K562 cells. CRISPR screen data (mean signal $log_2$FC, N=2) and sgRNA locations (bars) for CRISPRi-HCR-FlowFISH (FF) (orange) and Tycko et. al. 2019 CRISPRi-growth (blue). **B**) Upset showing number of overlapping PAM coordinates across 5 screens in *GATA1* and **C**) *MYC* (red: CRISPRi growth screen, blue: CRISPRi HCR-FlowFISH) **D**) percentage of exon (grey, total N=3642, 1739, 1735 from left to right) or K562 DHS targeting guides (black, total N=3472, 1570, 1592 from left to right) with significantly high $log_2$FC effect sizes (Z-score P value<0.001).

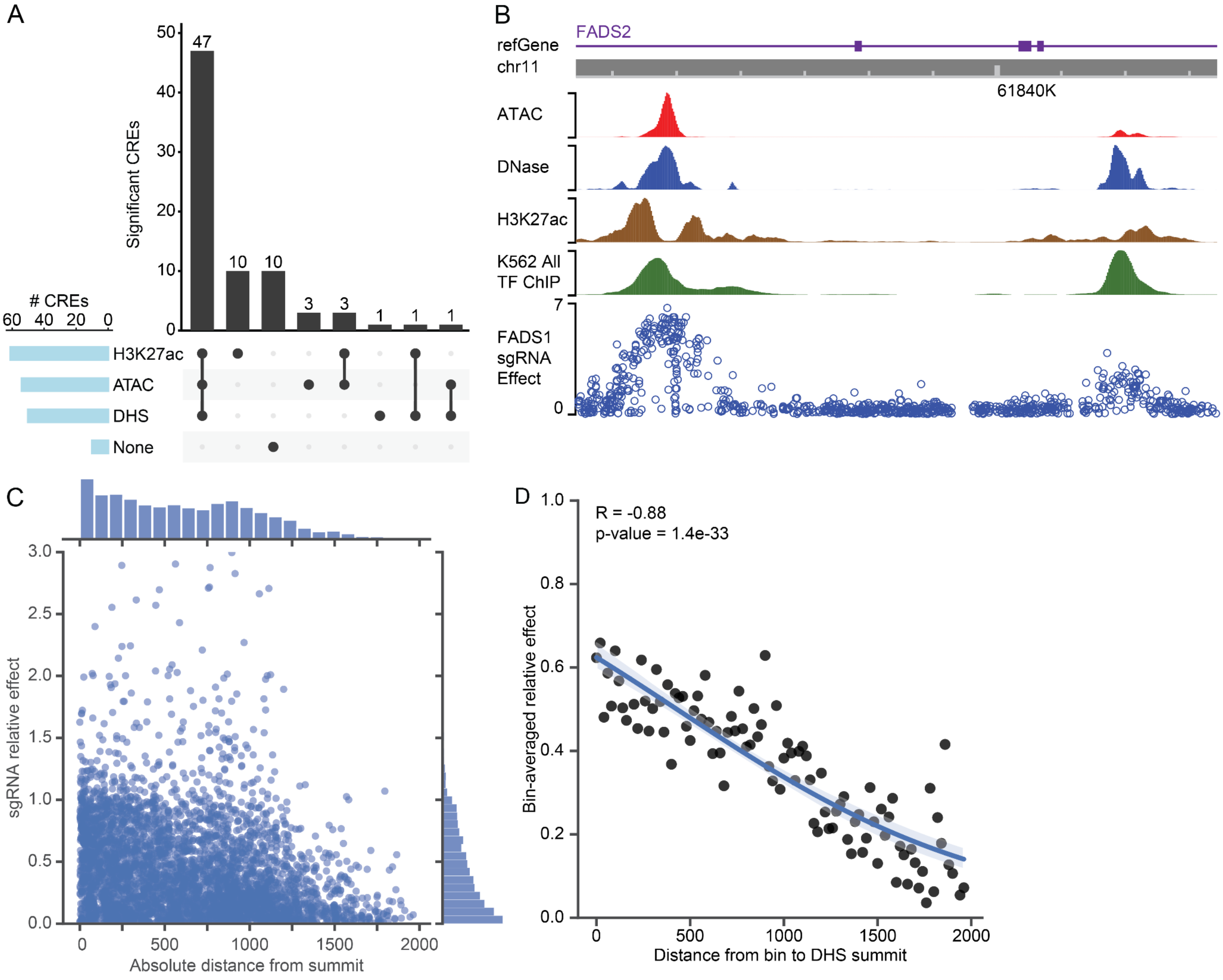


**Supplementary Fig. 4. Selecting cCREs and targeting sgRNAs near DHS summits**

**A**) Epigenetic feature peak intersections with significant CREs identified in 16 HCR-FlowFISH screens. **B**) Browser track highlighting two significant enhancers within *FADS2*. The K562 All TF ChIP track was created by concatenating all ENCODE K562 TF ChIP-seq experiments, and de-duplicating non-unique peak calls. The height of the track represents the number of unique TFs with peaks at a position. The average effects of each sgRNA from the *FADS1* HCR-FlowFISH screen (n=2 replicates). **C**) The effects of all sgRNAs across all HCR-FlowFISH screens within 2000 bases of a significant enhancer's DHS peak are plotted, normalized to the average effect of all sgRNAs in their enhancer. **D**) Same as (**C**), except sgRNAs are separated into 20 bp bins, with the mean of the sgRNA's enhancer-relative effects plotted for each bin; loess regression line drawn in blue.

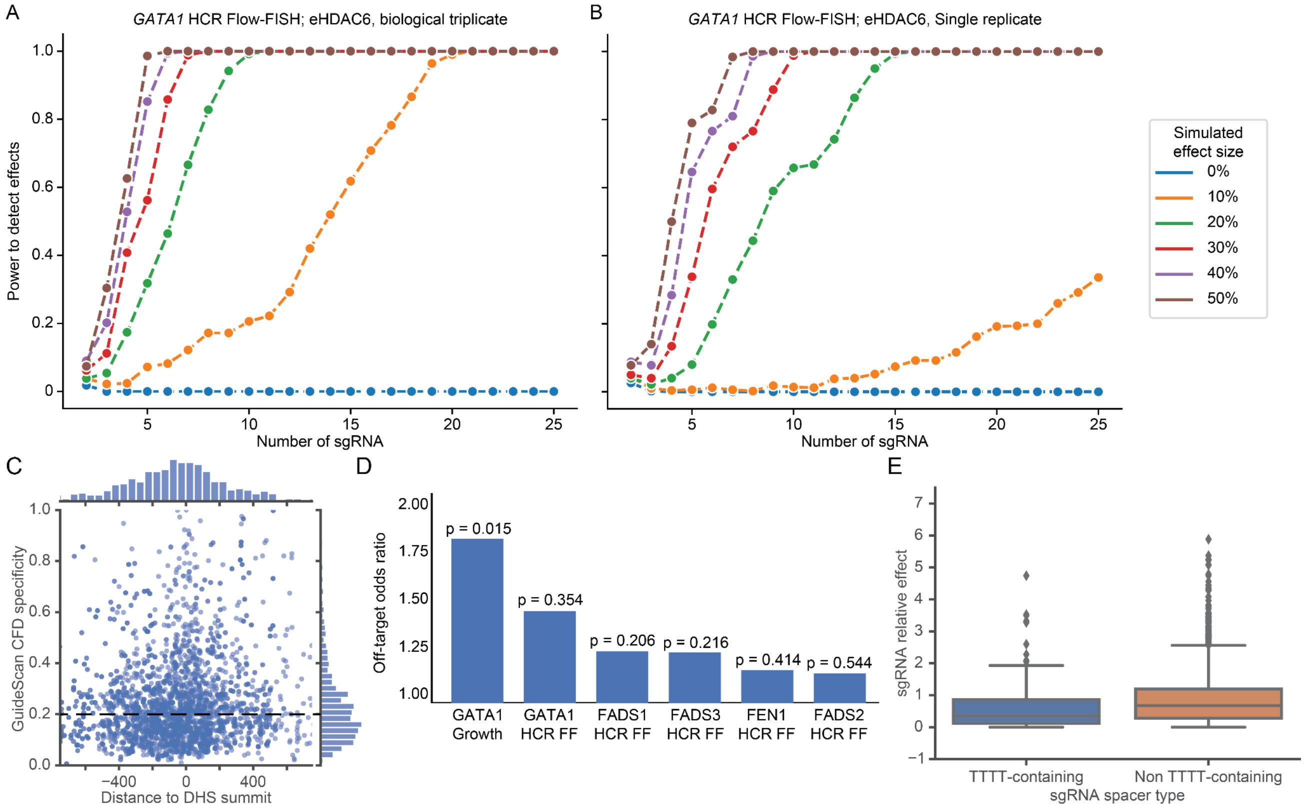


**Supplementary Fig. 5. Analysis of sgRNA specificity and power related to sgRNAs per element**

A) The power to detect significant effects on gene expression varies as a function of the number of sgRNAs targeting each element and the effect size of that element. Power was computed by simulations based on the average sgRNA effects from three biological replicates of GATA1 CRISPRi-FlowFISH data, where the individual sgRNA effects in the eHDAC6 element were scaled such that the average adjusted effect of all sgRNAs in the enhancer was 15%, 25%, or 35%. B) Same as (A), except for a single replicate. Note the loss of power with fewer replicates. C) Scatterplot of sgRNA PAM distance to DHS summits against their GuideScan CFD specificity scores, for all GuideScan sgRNAs in HCR-FlowFISH identified CREs that intersect DHS and H3K27ac peaks. Horizontal dashed line indicates GuideScan CFD specificity threshold of 0.2. D) Enrichment of sgRNAs with significant effects among sgRNAs with low specificity scores (GuideScan CFD<0.2) in regions at least 1 kilobase away from any DHS peak in K562 cells for the indicated screens. E) Distribution of sgRNA effects normalized to the average effect of all sgRNAs in their respective CREs, for sgRNAs with spacers that do or do not contain a ‘TTTT’ U6 termination sequence, using sgRNAs that target significant enhancers that intersect DHS and H3K27ac peaks. Boxes show the quartiles, with a line at the median, lines extend to 1.5 times the interquartile range, and dots show outliers. TTTT-containing sgRNA n=195; Non TTTT-containing sgRNA n=3940.

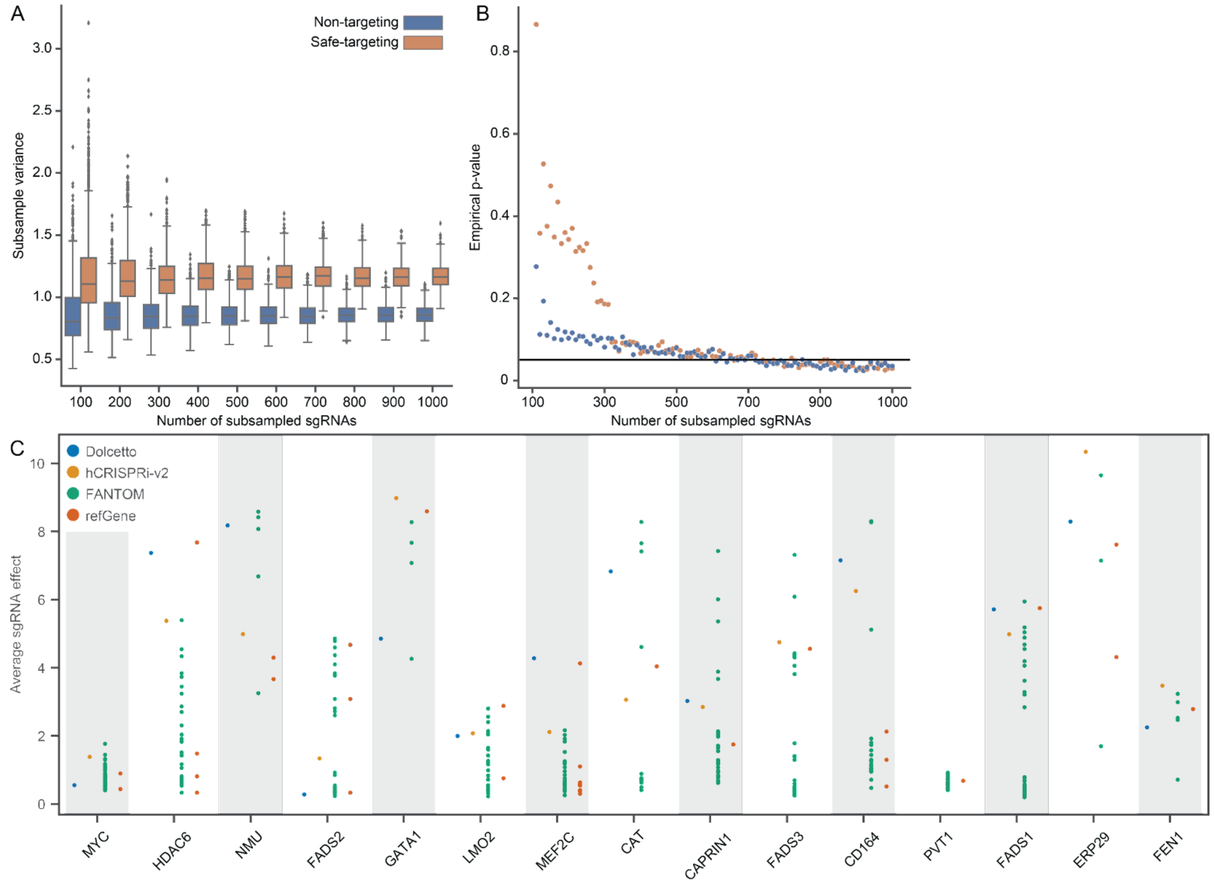


**Supplementary Fig. 6. Evaluating methods of selecting negative and positive control sgRNAs**

A) Boxplot of subsample variances for negative control sgRNAs in the CD164 HCR-FlowFISH screen, in increments of 100 sgRNAs subsampled 1000 times each from a total of 1000 sgRNAs for each type of negative control sgRNA. Each type of negative control was subsampled separately. Boxes show the quartiles, with a line at the median, lines extend to 1.5 times the interquartile range, and dots show outliers. B) Empirical P values from Levene's test on subsampled negative control sgRNAs, in increments of 10 sgRNAs subsampled 1000 times, compared to the entire set of the respective type of negative control sgRNA. P=0.05 threshold is indicated by the black line. C) Comparison of the average effect from both biological replicates of the 10 sgRNAs closest to the FANTOM5- and refGene-nominated TSSs for the HCR-FlowFISH genes against the sgRNAs provided by the Dolcetto or the hCRISPRi-v2 libraries, which may target one or more of these–or distinct–TSSs. Each point reflects an individual TSS (for the FANTOM5 and refGene TSSs) or the set of 4-10 sgRNAs from the Dolcetto or hCRISPRi-v2 libraries that were tested in the HCR-FlowFISH screens.

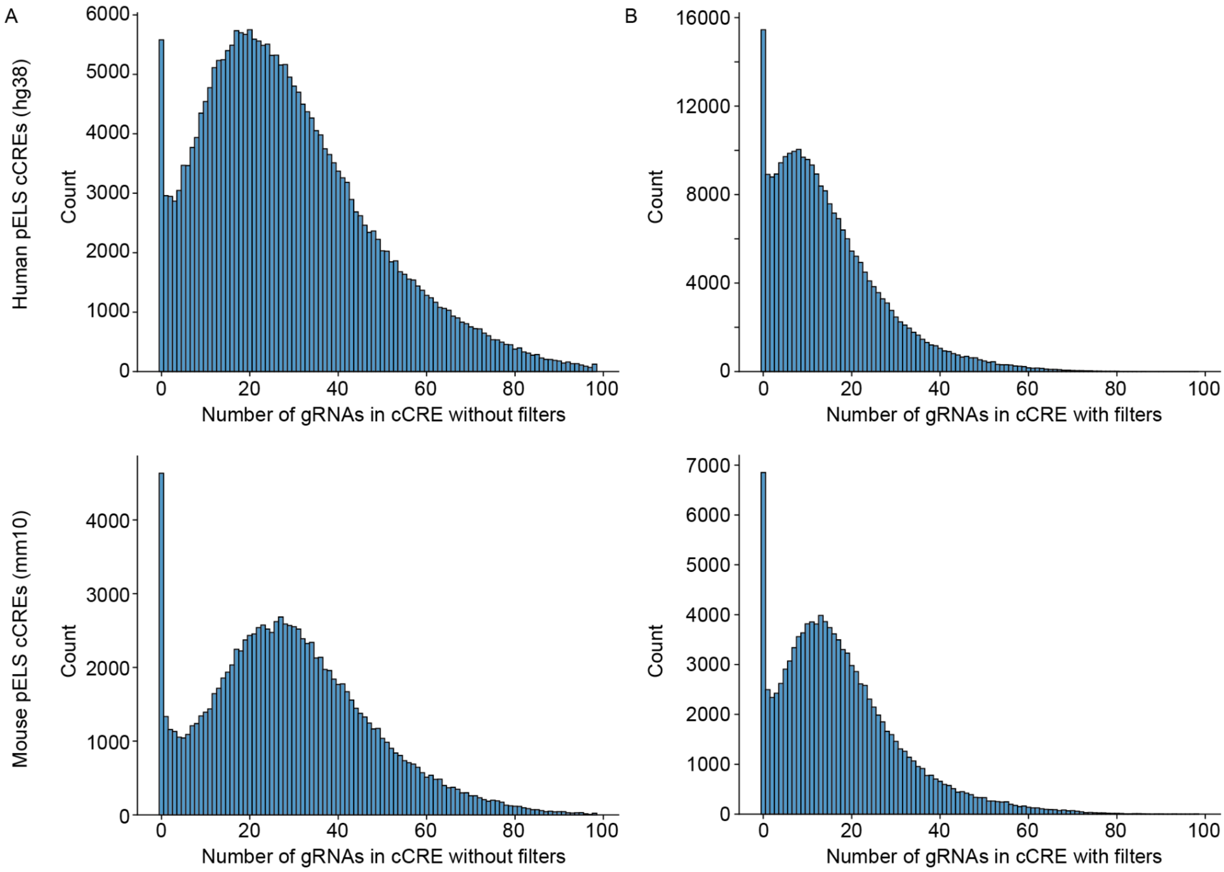


**Supplementary Fig. 7. GuideScan2-designed sgRNAs targeting all cCREs from the ENCODE SCREEN portal.**

**A**) The distribution of GuideScan2-designed sgRNAs per cCRE in the ENCODE SCREEN Registry v4 proximal enhancer-like signature (pELS) cCRE sets for human (top) and mouse (bottom) without and **B**) with filters to remove sgRNAs with a 'TTTT' sequence or GuideScan2-aggregated CFD specificity score<0.2.

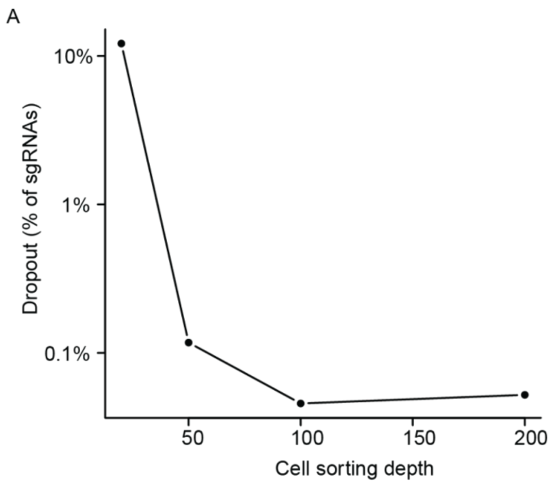


**Supplementary Fig. 8: Guide drop out rate with varying HCR-FlowFISH sorting depths**

**A**) sgRNA dropout rates (sgRNA with <10 mapped reads for low- or high-expression sorting bins) for varying cell sorting depth (20x, 50x, 100x, 200x) for CRISPRi *GATA1* HCR-FlowFISH performed at 2000x sequencing depth (n=1 replicate).

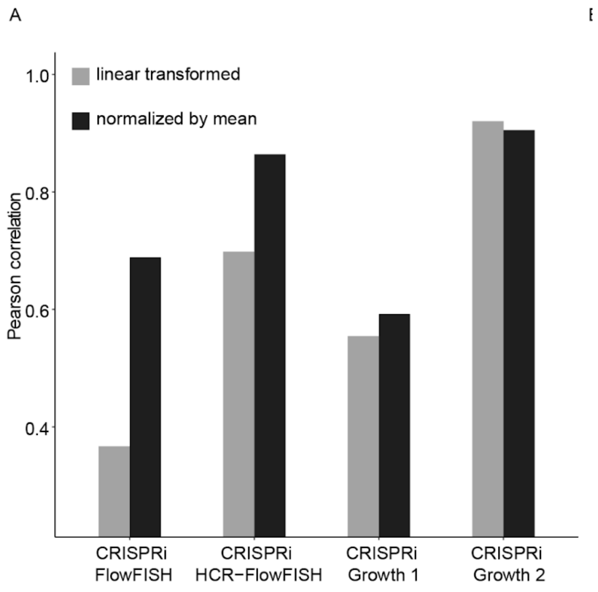


**Supplementary Fig. 9: Comparison of effect size normalization methods**

**A**) Comparison of linear transformed and mean-normalized ( $log_2((1 + (A_i\ /mean(A)))/(1 + (B_i\ /mean(B\ ))))$ ) effect size calculation by Guide-wise Pearson correlation of $\log_2$FC between two bio-replicates using different normalization methods (using ~250 sgRNAs commonly used across the GATA1 screens)

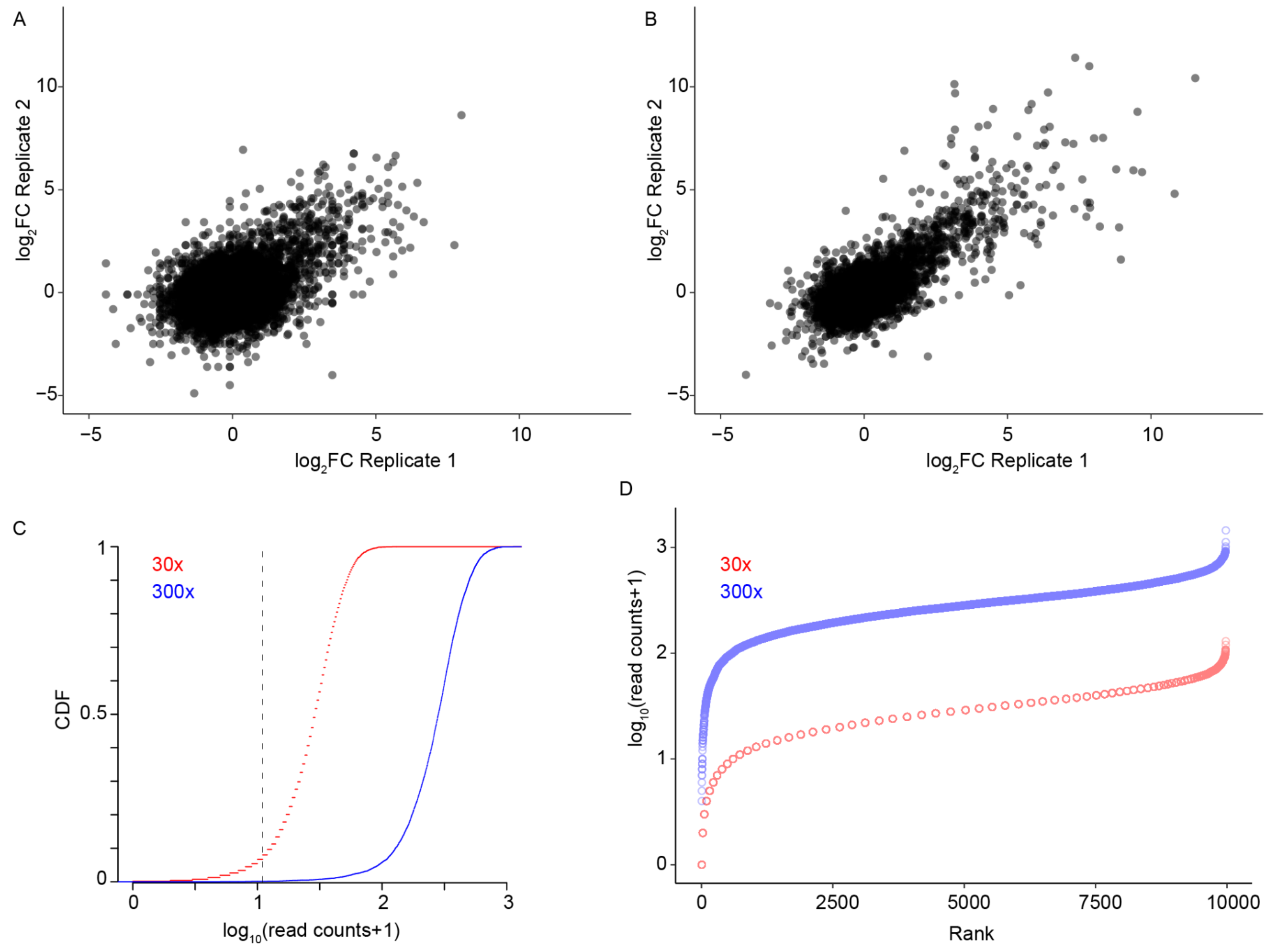


**Supplementary Fig. 10: Representative bootstrap samples for low and high sequencing depths using K562 GATA1 locus CRISPRi growth screen**

**A**) Biological replicate 1 $\log_2$FC vs Biological replicate 2 $\log_2$FC (Z-score) for 30x bootstrapped sequencing depth (9977 sgRNAs, R=0.45). **B**) Biological replicate 1 $\log_2$FC vs Biological replicate 2 $\log_2$FC for 300x (R=0.73). **C)** Empirical cumulative distribution function of $\log_{10}$(sgRNA read counts) **D**) Dropout plot (rank of sgRNA read counts vs $\log_{10}$(1+read counts) at 30x (red) and 300x (blue) bootstrapped sequencing depth.

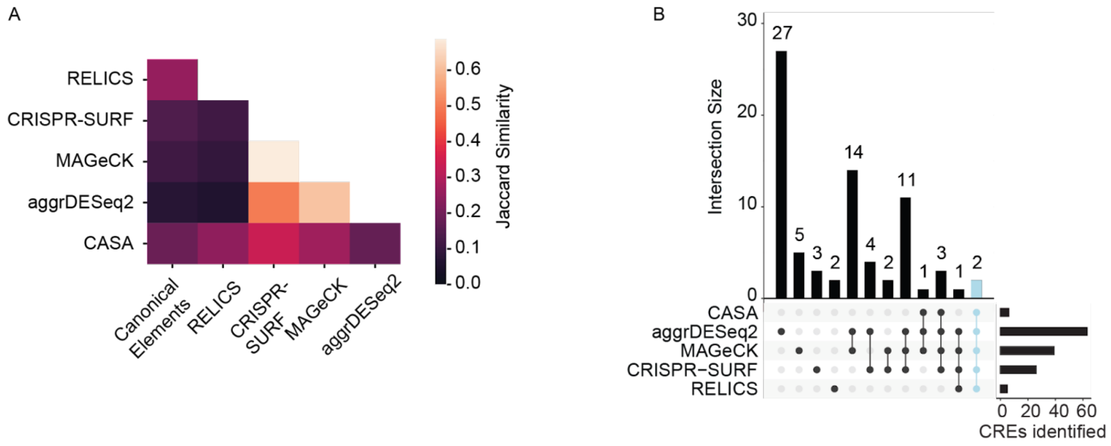


**Supplementary Fig. 11. Overlap between peak calls on specificity-filtered CRISPRi tiling screen of *GATA1* locus**

**A**) Quantification of pairwise overlap of peak calls by Jaccard Similarity. High values (lighter colors) correspond to greater overlap. **B**) Upset plot of intersections of peaks identified by the 5 peak callers.

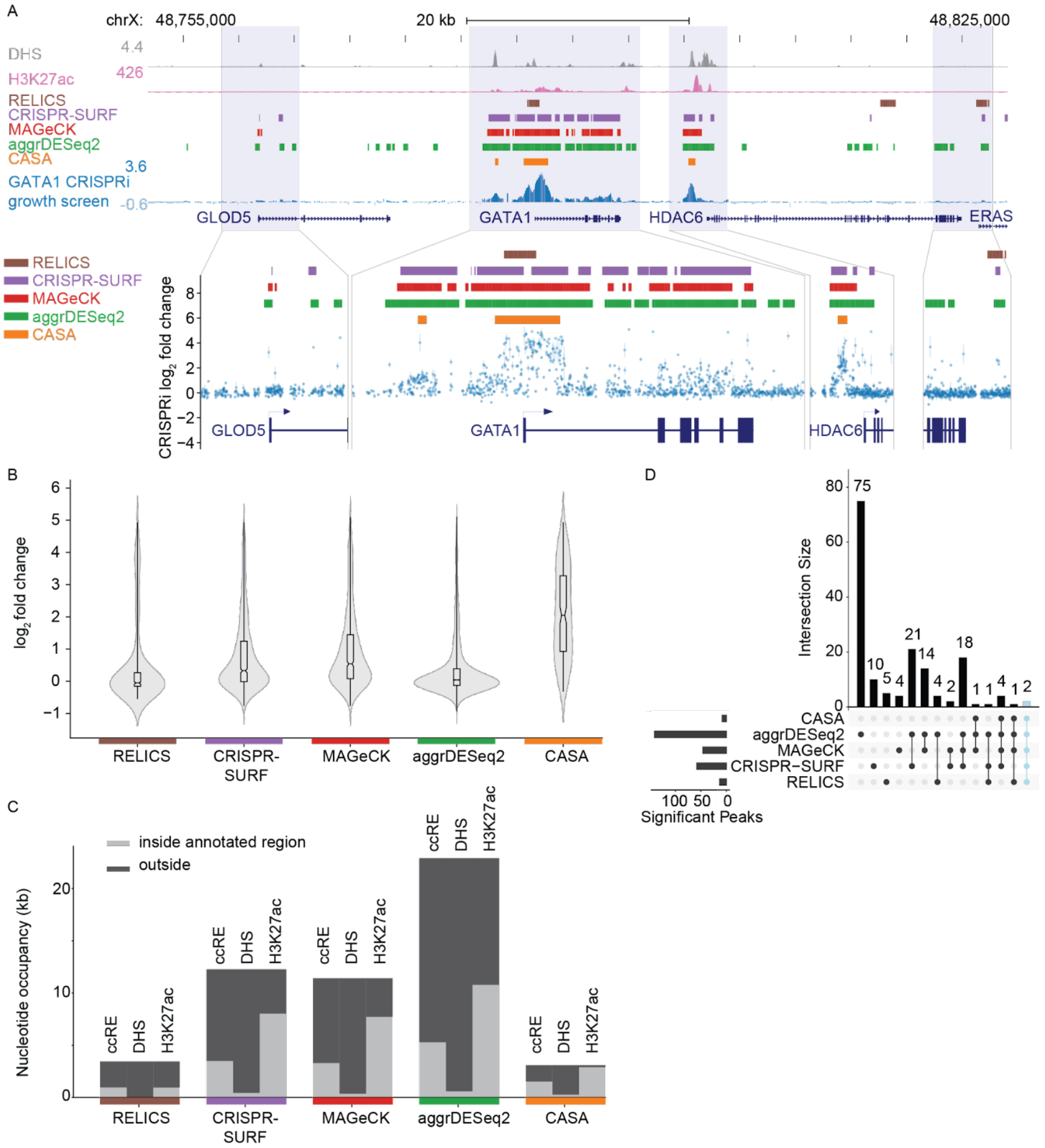


**Supplementary Fig. 12. Peak calls without filtering out low specificity sgRNAs**

**A**) sgRNA mediated growth effects (blue), H3K27ac-ChIP signal (pink), and DNase Hypersensitivity signal (gray) for a CRISPRi growth screen at the GATA1 locus, without removing low specificity sgRNAs. Dense tracks show peak calls using 5 different CRISPR screen analysis tools: CASA (orange), aggrDESeq2 (green), MAGeCK (red), CRISPR-SURF (purple), and RELICS (brown). Zoomed-in regions show individual sgRNA effects (points, mean; bars, min-max range of observations between n=2 replicates). **B**) Distribution of average sgRNA effects from two experimental replicates for sgRNAs falling within peaks identified by different CRISPR screen analysis tools (center line, median; notch confidence interval of the median; box limits, first and third quartiles; whiskers range of all data points; violin, kernel density estimation). **C**) Total peak area inside (light gray) and outside (dark gray) of annotated chromatin features for each peak caller. **D**) Intersections of peaks identified by the 5 peak callers.

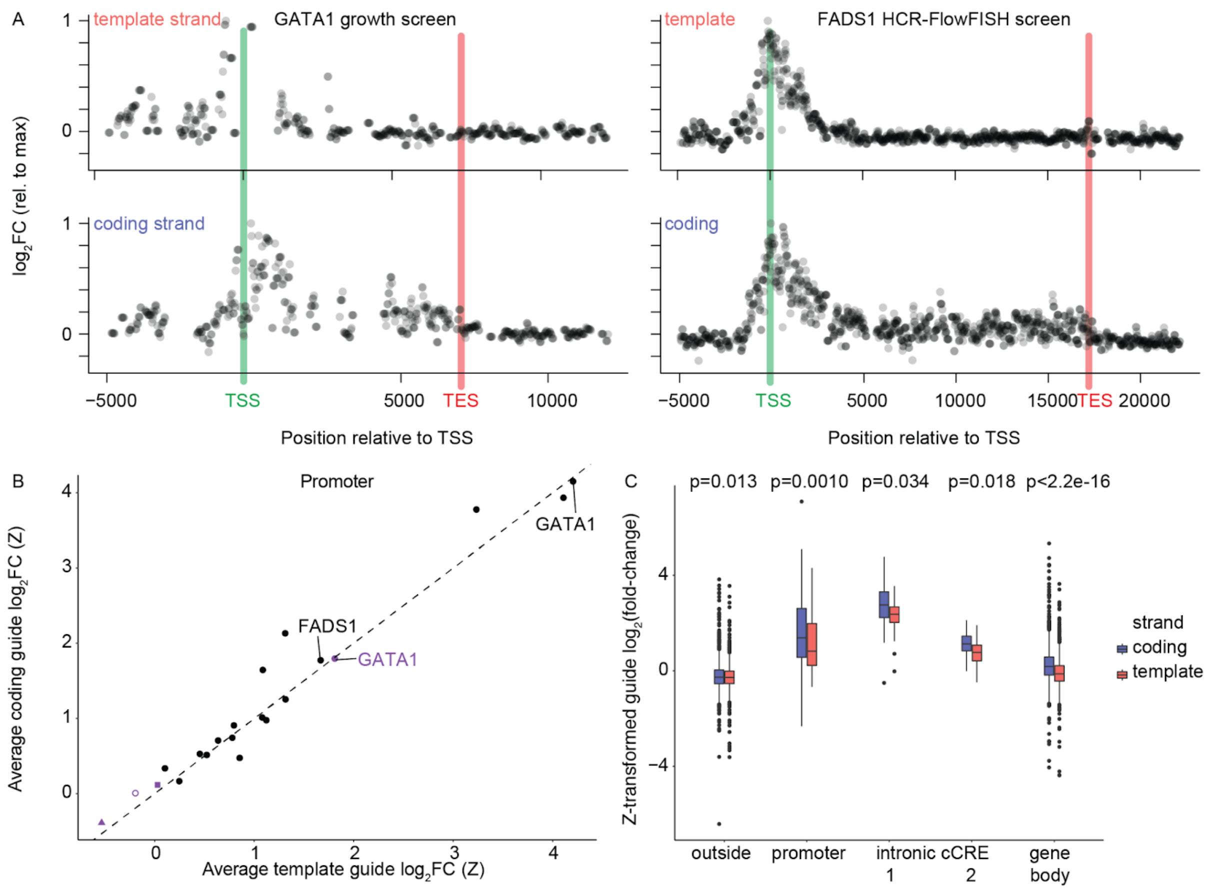


**Supplementary Fig. 13. CRISPRi strand bias in the gene body**

**A**) CRISPRi tiling screen effects shown relative to the position of the TSS and transcription end site (TES). The TES is defined as the end of the transcript in UCSC RefGene (hg38). Each point shows the average normalized sgRNA effect from two screen replicates. **B**) Strand-specificity across screens tiling 19 loci for sgRNAs targeting the promoter. Each point is the average effect of all sgRNAs targeting that region, averaged across two biological replicates, with color indicating the phenotypic readout (e.g. growth), and shape indicating the type of CRISPR perturbation used (e.g. CRISPRi). **C**) Distribution of sgRNA effects in a CRISPRi FlowFISH tiling screen for FADS2 regulatory elements. Outside refers to sgRNAs that are outside the gene body, promoter, and K562 DHS peaks but are within 1 Mb of the FADS2 TSS. The two intronic CREs are annotated in **Fig. 6B**. T-test P values for the comparison across strand categories are shown. Boxes show the quartiles, with a line at the median, lines extend to 1.5 times the interquartile range, and dots show outliers (left to right: N=2105, N=1935, N=107, N=126, N=32, N=26, N=27, N=19, N=1940, N=1786).

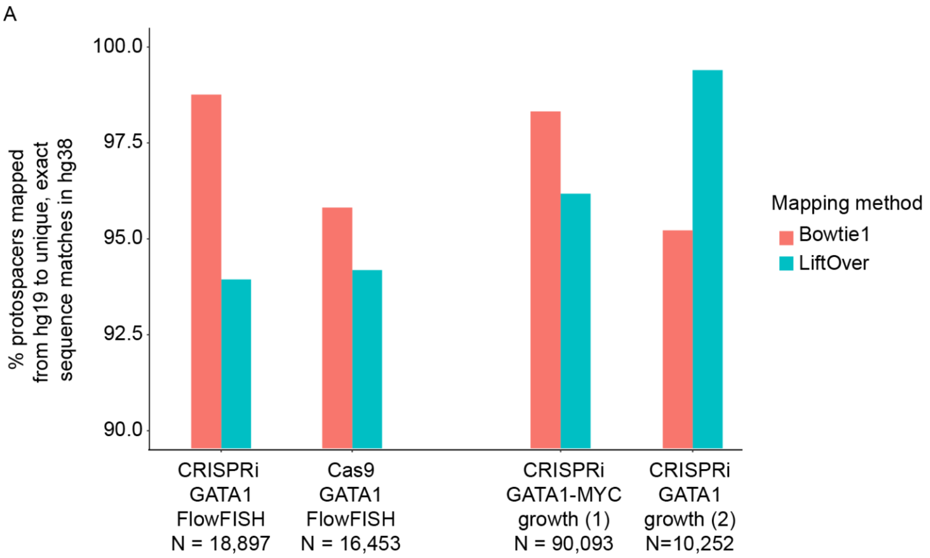


**Supplementary Fig. 14. Mapping sgRNAs to reference genome for data standardization**

**A**) Four CRISPR screen sgRNA libraries in hg19 were mapped to hg38 using bowtie1 and LiftOver. For three out of the four libraries, bowtie1 resulted in higher rates of unique and exact sequence mapping (**Supplementary Section 6**). CRISPRi-Growth datasets are (1) Tycko *et. al.* 2019 and (2) Fulco *et. al.* 2019.

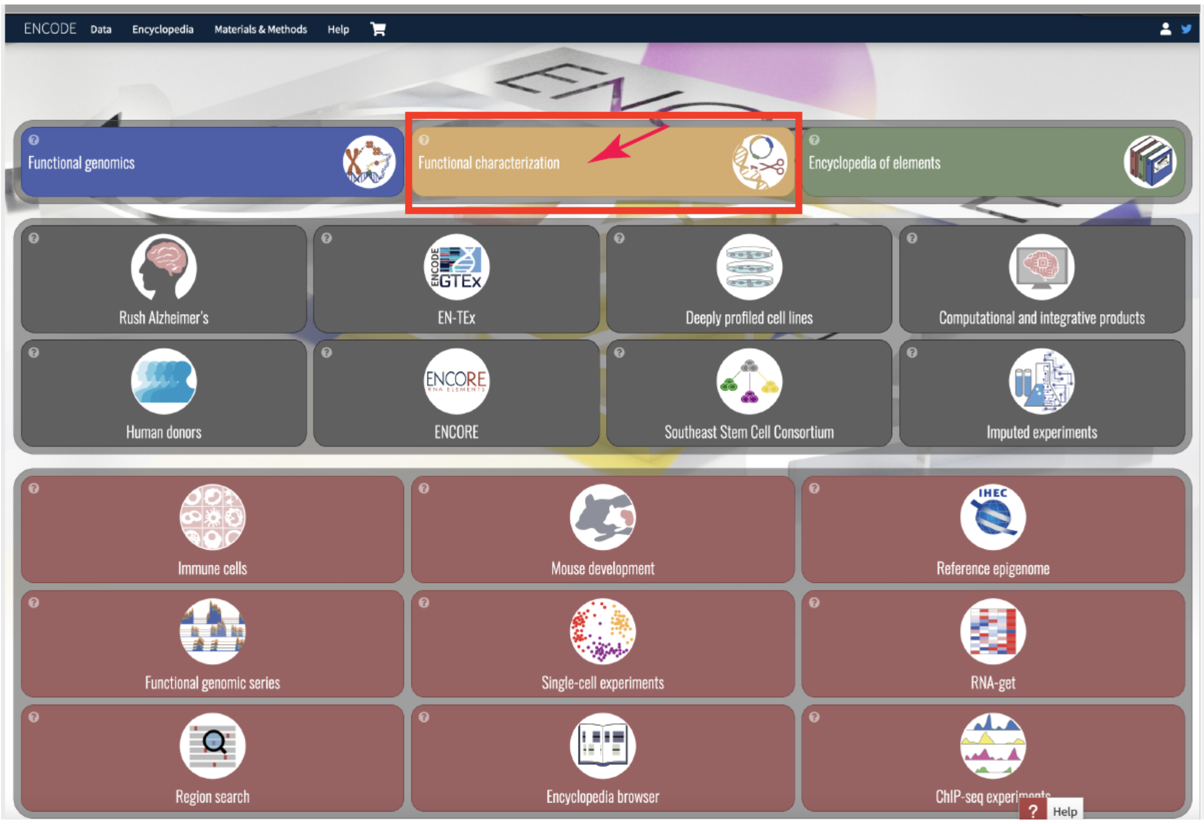


**Supplementary Fig. 15. 'Functional Characterization' card on the ENCODE home page**

Representative image of the ENCODE portal home page. The red box indicates the card to select to navigate to the Functional Characterization assay datasets.

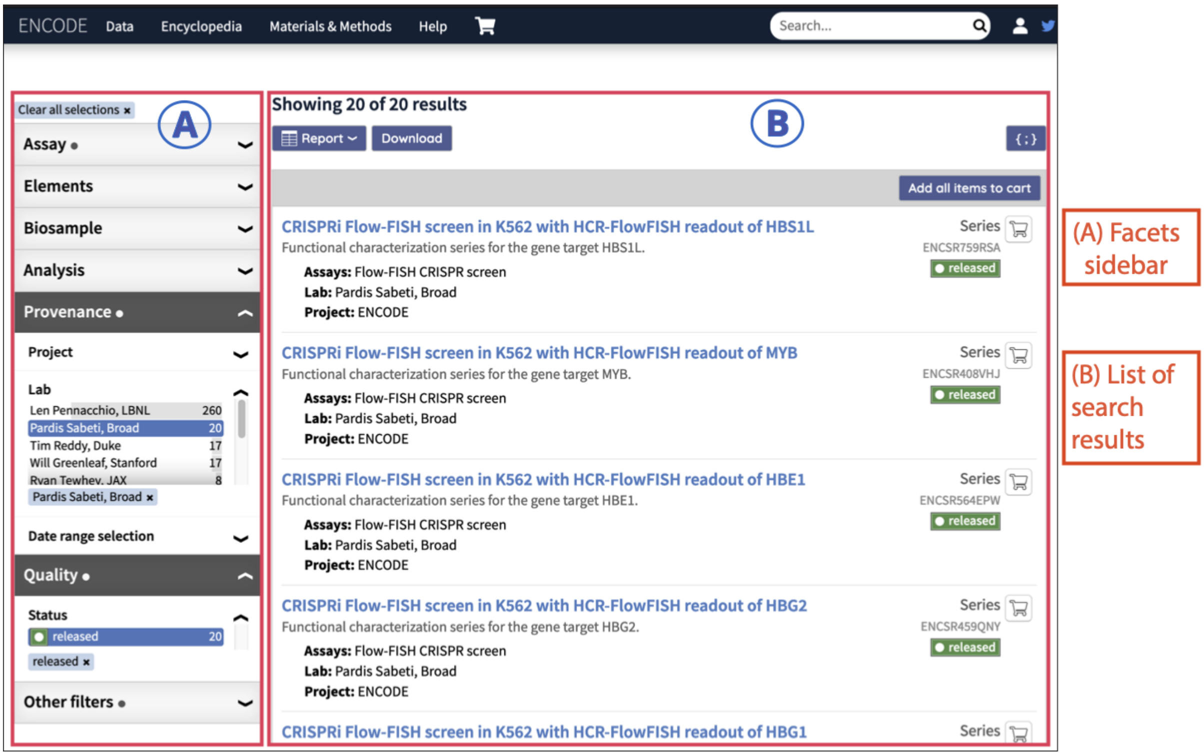


**Supplementary Fig. 16. Filtering search results in the ENCODE portal**

**A**) A view of the facets sidebar with the items that should be selected and the list of the resulting ENCODE Functional Characterization Series. **B**) Each Series is shown with a brief summary of the biological material, assay name, and a link to its individual experiment series summary page with more metadata details. The second Series from the list (ENCSR408VHJ) is selected as an example and further exploration.

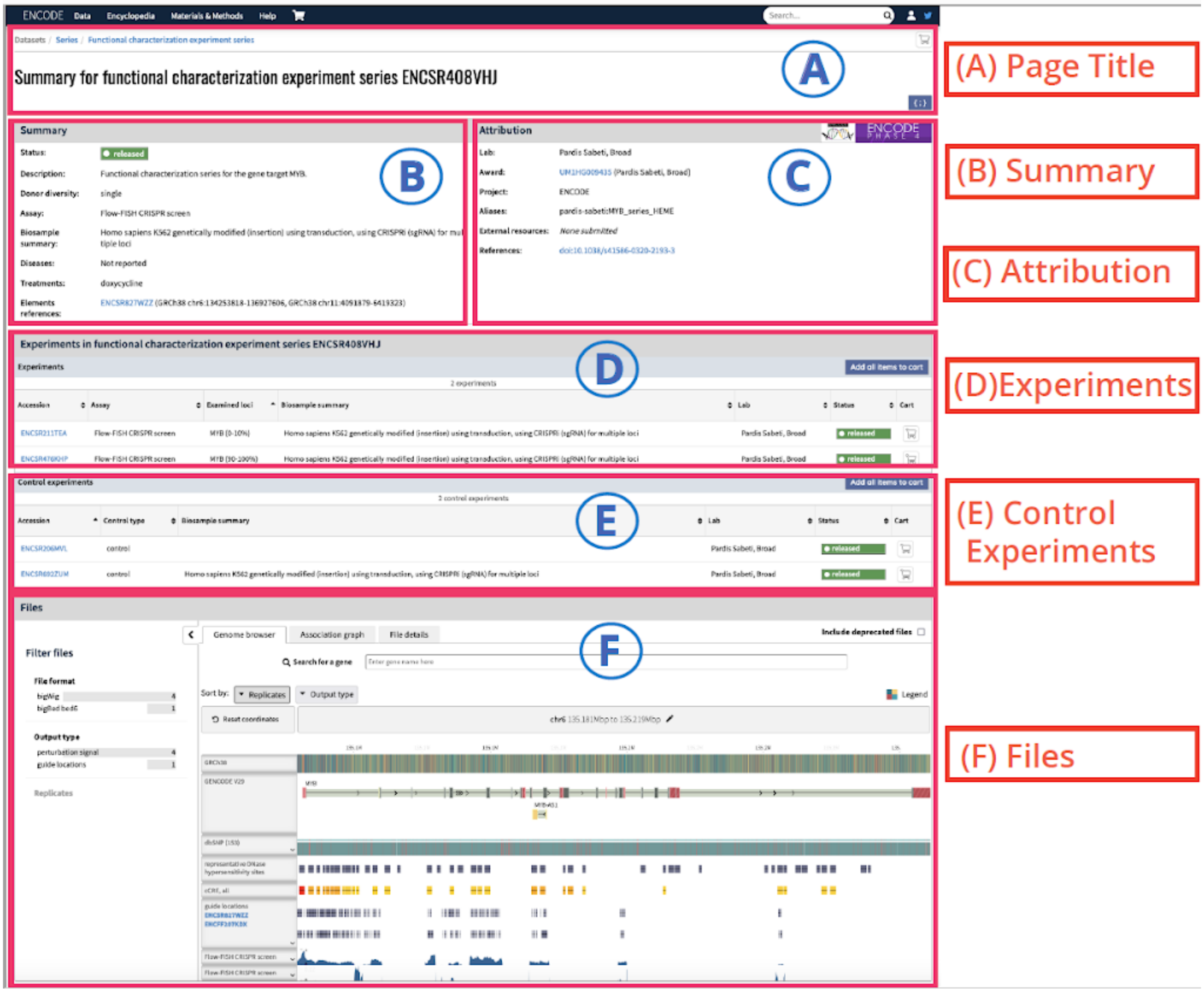


**Supplementary Fig. 17. An experiment series summary page**

The page is organized into six distinct sections. **A**) Page Title: providing the series title and accession. **B**) Summary: providing key information about the series. **C**) Attribution: providing information about the lab that performed the experiment, aliases and external resources. **D**) Experiments: providing information about all experiments that have been included in the series. **E**) Control Experiments: providing information about all auxiliary and control experiments that have been included in the series. **F**) Files: contains information about the files associated with the series.

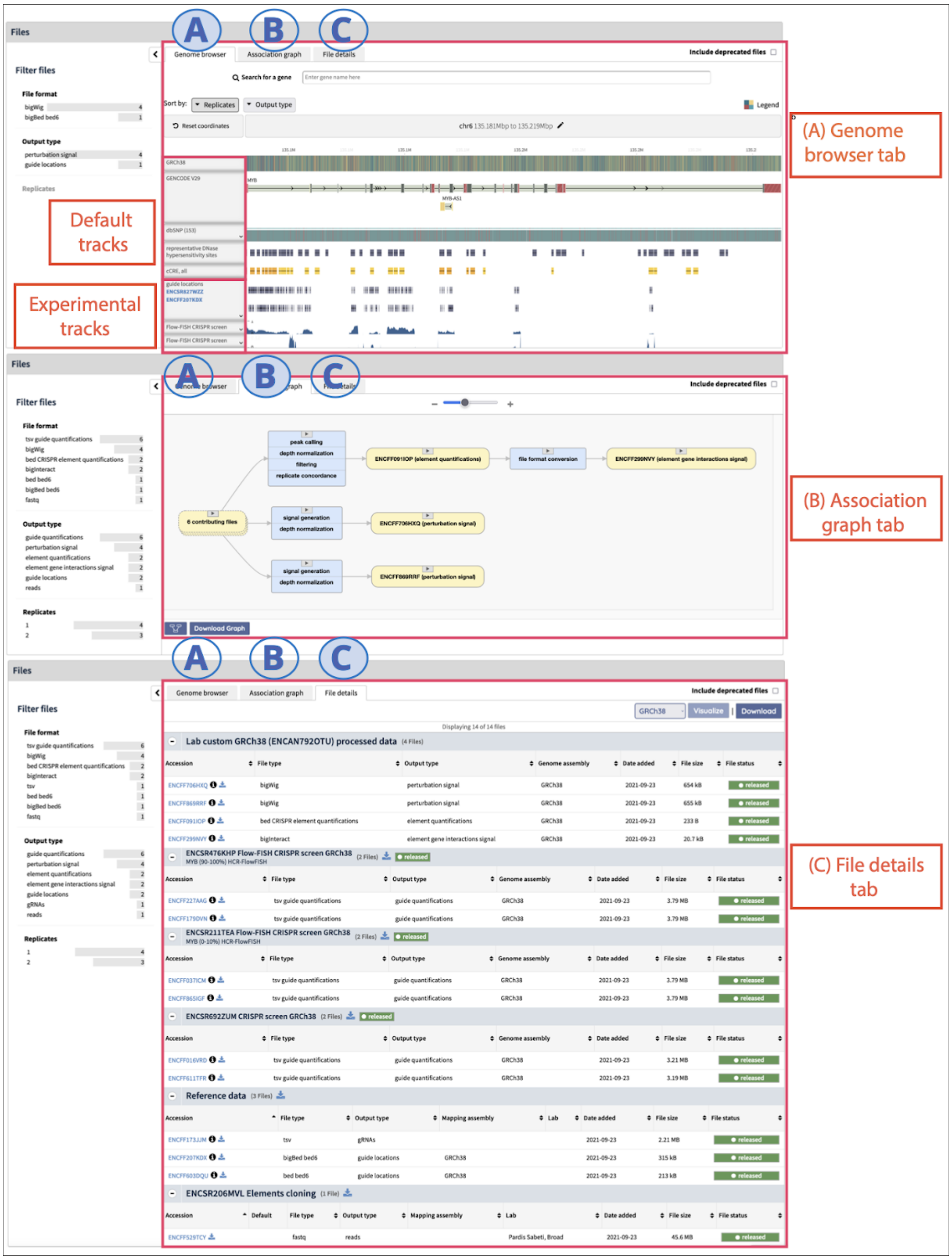


**Supplementary Fig. 18. The 'Files Section' of the experiment series summary page**

The Files Section of the experiment series summary page contains three tabs. **A**) Genome browser tab: visualizes tracks using the embedded Valis genome browser. **B**) Association graph tab: displays the data provenance and derivation of downstream processed files. **C**) File details tab: lists the files that are associated with the series.

#### Supplementary Tables

A description of the contents of all Supplementary Tables is provided below. Supplementary Tables 1-7 and 9-17 are provided in one separate file. Due to size, supplementary Table 18 is provided as a separate file. Due to size, Supplementary Table 8 is provided as a separate file and can be accessed at: https://drive.google.com/drive/folders/1vjyPEmGVmmV74AJNA9UxuathYR0BijJo

Supplementary Table 1: CRISPR screens in the ENCODE data portal.
Supplementary Table 2: CREs identified from available CRISPR screens performed in human biosamples.
Supplementary Table 3: Overview of noncoding CRISPR screen approaches.
Supplementary Table 4: Annotation files used in K562 meta analysis.
Supplementary Table 5: CREs identified from available CRISPR screens performed in K562s.
Supplementary Table 6: Fisher's exact test results for enrichment of cell-type agnostic and K562 annotations in functional regulatory elements.
Supplementary Table 7: Signal values and test results for comparison of K562 feature signal in significant CREs versus perturbed regions.
Supplementary Table 8: Genome-wide ENCODE SCREEN cCRE GuideScan sgRNA libraries.
Supplementary Table 9: Negative control sgRNAs used in this study.
Supplementary Table 10: Summary of common sgRNA design tools.
Supplementary Table 11: Summary of analysis tools used in peak calling comparison.
Supplementary Table 12: *GATA1* locus sgRNAs for validations.
Supplementary Table 13: CRISPR screen guide_quantification file format and contents.
Supplementary Table 14: CRISPR screen element_quantification file format and contents.
Supplementary Table 15: Accession IDs from ENCODE portal used in this study.
Supplementary Table 16: hg38 Safe-targeting sgRNA library.
Supplementary Table 17: GuideScan2 sgRNA counts at SCREEN cCREs.
Supplementary Table 18: sgRNA library and read counts for *GATA1* titration experiments.

### Materials and Methods

#### Cell lines and cell culture

K562 cells with a doxycycline-inducible CRISPRi-BFP were a gift of the Lander lab and identical to those used in previous studies[5]. In that study, the cells were generated by (i) transducing K562 cells with a construct expressing rtTA linked by IRES to a neomycin resistance cassette expressed from an EF1α promoter (ClonTech, Mountain View, CA) and selecting with 200 μg/mL G418 (Thermo Fisher), then (ii) transducing these rtTA-expressing K562 cells with a KRAB-dCas9 construct. Cells expressing BFP were selected by fluorescence activated cell sorting (FACS). Cells were grown in RPMI 1640 GluteMAX (Gibco) with 10% heat inactivated FBS (HI-FBS, Gibco).

#### GATA1 screen with varied cell coverage

A previously described non-coding *GATA1* lentiviral library was used[5]. CRISPRi-BFP was induced for 24 h with a final concentration of 1 μg/ml doxycycline (VWR). Active CRISPRi was checked by confirming dox-induced BFP signal was observed in >90% of cells by flow cytometry (Sony MA900). Cells were grown for 2 weeks post transfection, following the HCR-flowFISH protocol exactly as previously described[5]. High and low expression bins (top and bottom 10% each), were also gated following previous HCR-FlowFISH protocol[5]. Cells were sorted at multiple folds of library size (25x, 50x, 100x, and 200x).

#### The ENCODE CRISPR Screen Database and overlap with cCREs

*Calculation of total perturbations and percent of genome perturbed*

Individual sgRNAs were aggregated across fully released experiments with sgRNA-level and/or element-level quantification files performed in human cell lines using the April 2022 data release (excluding single cell gene expression readouts). Note that three experiments in the April 2022 data release were removed in the August 2022 data release (**Supplementary Table 1**). These experiments have been re-released as of November 2022 and are included in "Experiments", "Biosamples", and "Genes/phenotypes" in **Figure 1B** but were excluded from the calculation of total perturbations, the calculation of percent of genome perturbed, and the significant CRE intersection. The coordinates of each sgRNA were adjusted based on the type of perturbation used in the corresponding experiment (Cas9 cutting: ±10 bp of PAM, dCas9: ±10 bp of PAM, dCas9-KRAB: ±150 bp of PAM) and lifted from hg19 to hg38 genome builds when necessary. The total number of perturbations was defined as the number of unique protospacer and coordinate combinations. These perturbation regions were then intersected with 100bp tiled bins across each chromosome, followed by merging of overlapping bins (`bedtools merge -d 1`), and the percent of the human genome perturbed was calculated by dividing the sum of bases within the tiled bins by the effective genome size (3,088,269,832 bp). The significant CREs from each experiment (defined by the contributing lab) were intersected with the same 100 bp tiled bins and similarly merged to generate the final CRE set (**Supplementary Table 2**).

#### K562 screens integrated analysis

*Experiments included in analysis*

ENCODE accession numbers and other experiment metadata are provided in **Supplementary Table 1**.

*Annotation data*

The genomic and epigenomic annotation files used in this analysis are provided in **Supplementary Table 4**.

*Aggregating screen perturbations and significant CREs*

Individual sgRNAs were aggregated across released experiments performed in K562s with sgRNA-level and/or element-level quantification files (April 2022 data release, excluding single cell gene expression readouts; **Supplementary Table 1**, 'Included_in_k562_meta'). The coordinates of each sgRNA were adjusted based on the type of perturbation used in the corresponding experiment as described above and lifted from hg19 to hg38 genome builds when necessary. These perturbation regions and the CREs from each experiment (defined by the contributing lab) were then intersected with 100bp tiled bins as described above to generate the perturbed and CRE sets, respectively. The CRE coordinates and feature overlap are provided in **Supplementary Table 5.**

*Enrichment testing and signal comparison*

The perturbed regions and CREs were intersected with the significant peak calls or predicted ENCODE SCREEN cCREs ("features"). Fisher's exact test was performed, comparing the number of features overlapping a CRE to the total number of features perturbed. To compare the signal of each feature between perturbed region and CREs, bigWig files were converted to bedgraph format using the UCSC utility 'bigWigToBedGraph'. Next, the perturbed regions and CREs were intersected with the bedgraph files containing fold change over background signal ("signal"). Signal values were then normalized by dividing by the element size, and a Student's t-test was performed, comparing the mean signal for each feature between perturbed, not significant regions, and CREs. Student's t-test and Wilcoxon test results, and mean and median signal values, with and without the element size normalization are reported in **Supplementary Table 7**.

##### CRISPR screen comparisons with individual sgRNA validations

sgRNA abundance and element activity values from CRISPR screens, and results from experimental validations, were obtained from supplemental materials for each of the cited publications. Pearson correlation values and associated P values between the validation assay and the screen result were calculated using the 'stat_cor' function from the R package 'ggpubr'.

##### Cross-screen analysis at *GATA1* and *MYC*

hg38 PAM coordinates were used to uniformly analyze and compare the five CRISPR screens from various labs. For screens with hg19 coordinates, their protospacer coordinates were first mapped to hg38 using bowtie1 using "-n --best" options (**Supplementary Section 6**). Then, the hg38 PAM coordinates for each screen were extracted by taking the three base pairs downstream of each protospacer, which were confirmed to contain the expected NGG sequence. For the *GATA1* locus, 250 such PAM coordinates were found to be shared across the five screens, and these common PAM coordinates were filtered out for their sgRNA GuideScan target specificity (>0.2), leading to 176 PAM coordinates which were used for pairwise effect size comparison of the five screens (**Fig. 1D,E** and **Supplementary Fig. 2B**). Effect sizes were computed using mean normalized $\log_2$FC (**eq 3.1,** provided below in **'Cell coverage / sorting depth titration experiments for HCR-FlowFISH'**). To compare CRISPR-Cas9 and CRISPRi's effects at exons and DHS, we obtained subsets of sgRNAs with significantly high $\log_2$FC effect sizes (Z-score P value<0.001). Then, we extracted significant sgRNAs that target exons or K562 DHS by overlapping their PAM coordinates with Ensembl-annotated exons and K562 DHS obtained by extending K562 DHS narrow peaks (ENCFF899KXH) by 500 base pairs in both directions from their centers. (**Supplementary Fig. 2D**).

##### Evaluating sgRNA effects in DHS or H3K27ac peaks

Significant, non-TSS-overlapping distal enhancer elements identified in any of the HCR-FlowFish screens that intersect both a DHS and H3K27ac peak were first selected. For each enhancer element, we calculated the mean effect of all sgRNAs within its intersecting DHS or H3K27ac peak region. The sgRNA intersections used the sgRNA's 3-base PAM coordinate window.

##### Evaluating sgRNA effects as a function of distance from the DHS summit

Significant, non-TSS-overlapping distal enhancer elements identified in any of the HCR-FlowFish screens that intersect both a DHS and H3K27ac peak were selected. We then selected all sgRNAs within 2 kb of the enhancer element's strongest intersecting DHS summit, and normalized their effect sizes to the mean of all sgRNAs intersecting that DHS peak (using the sgRNA's 3-base PAM coordinate window).

For **Fig. 2B**, we took the sgRNA coordinate and expanded it by ±150 bp to conservatively approximate KRAB's repressive window, and assigned each base position that sgRNA's normalized effect size. If multiple expanded sgRNA windows overlap, then their effects are averaged per base position. This data was converted into a bigWig file, and we used deepTools to plot the distance-dependent sgRNA effects along with DNase-seq and H3K27ac ChIP-seq signal P value tracks. For **Supplementary Fig. 4C**, we plotted the normalized effects of all sgRNAs in the same set of significant enhancers as above, as a function of distance to their respective enhancer's DHS center. In **Supplementary Fig. 4D**, normalized sgRNA effects are averaged into 20-base bin intervals.

##### Effect size-dependent sgRNA number per element power analysis

For the guide downsampling analysis, we took guide-level effect sizes from the CRISPRi-FlowFISH screens targeting the *GATA1* locus and averaged the effect sizes from two biological replicates. We then took the sgRNAs targeting the e-GATA1 enhancer and rescaled their effects so that the average of all 37 sgRNAs was either a 0-50% perturbation, in steps of 10%, of *GATA1* expression. For each number N of sgRNAs, we sampled N sgRNAs from the scaled distribution, computed a Welch's T-test P value (`equal_var=False, dof=1`) against all non-targeting negative control sgRNAs, performed a BH-correction with all elements tested in the screen, and tested for FDR<0.05. We repeated this procedure 500 times for each (effect size, guide number) pair and computed power as the fraction of times we correctly rejected the null hypothesis.

##### Off-target sgRNA enrichment analysis

For each respective screen, we selected sgRNAs located at least 1 kb away from any DHS peak, regardless of significance, or significant element. We used GuideScan to obtain sgRNA aggregated CFD scores, a summary score of off-target specificity based on the weighted likelihood of off-target activity across a full list of potential off-target sites, and separated sgRNAs into low-specificity (CFD<0.2) or high-specificity (CFD≥0.2). We then calculated the proportion of sgRNAs in each specificity category that had effect sizes more than two times the standard deviation of negative controls from the mean of the negative controls, and performed a Fisher's Exact Test to derive a P value for each odds ratio.

##### Safe vs. non-targeting negative control variance statistical analysis

For **Supplementary Fig. 7**, negative control sgRNAs were subsampled 1,000 times each, in increasing increments of 10 sgRNAs. For each subsample, we performed Levene's test against the full set of 1,000 of the respective type of negative control sgRNAs. We then calculated the percentage of times that the result of Levene's test was significant (P<0.05) – that is the number of times variance between the subset and the whole set is statistically different – from the 1,000 subsamples for each increment. This percentage is the empirical P value, such that the black threshold line of P=0.05 means that out of 1,000 subsamples, only 50 had significantly different variances compared to the variance of the full set of that respective type of negative control sgRNA.

##### Promoter-targeting "positive control" sgRNA selection analysis

For **Supplementary Fig. 7C**, we selected all TSSs provided by the FANTOM5 database that pass a relaxed Timo TSS classification score of 0.14 for the genes measured by HCR-FlowFISH. We calculated the average effects of the 10 closest sgRNAs to each TSS position; where a TSS window is provided, we used the first transcribed base position to calculate absolute sgRNA distances. To compare these sgRNA against those provided by genome-wide CRISPRi libraries (Broad Dolcetto[9] and hCRISPRi-v2[10]), we selected the sgRNAs whose spacers match those tested in the HCR-FlowFISH screening libraries; the sgRNAs from hCRISPRi-v2 follow a G+19 base spacer convention, so the 5′-most base from the HCR-FlowFISH spacer sequences was trimmed to facilitate spacer sequence matching. Since these libraries often provided lower scores than the optimal TSS, we aimed to provide a heuristic method of selecting TSS-targeting sgRNAs by selecting the TSS with the greatest Pol-II ChIP-seq signal (TSS provided by RefGene, total Pol-II ChIP-seq signal was calculated in a window ±500 bp around the TSS), and picking the 10 nearest sgRNAs.

##### Cell coverage / sorting depth titration experiments for HCR-FlowFISH

HCR-FlowFISH experiments at *GATA1* were performed using guide libraries, K562 cell lines, transcript detection, sorting, and sequencing strategies as previously described[5]. To evaluate the effects of sampling cell numbers at different levels of complexity, defined as the number of observations per number of sgRNAs used, we performed two replicates of the *GATA1* library and partitioned them into different sorting depths. The same library was sorted into 20X, 50X, 100X, and 200X the guide library size. To assess the impact of sequencing complexity, each sorting strategy was sequenced at a depth of more than 2000X.

Effect size of each sgRNAs was computed using eq 3.1 to underweight sgRNAs with low read counts by normalizing read counts by their mean, which led to higher bio-replicate reproducibility relative to $\log_2$FC ($log_2((1 + A_i)/(1 + B_i))$) or its linearly transformed variants (e.g. Z-transformed $\log_2$FC and eq 3.2) (**Supplementary Fig. 9**):

$$log_2FC_i = log_2((1 + (A_i / mean(A)))/(1 + (B_i / mean(B)))) \quad \text{eq 3.1}$$

$$log_2FC_i = log_2(((1 + A_i) / sum(A))/((1 + B_i) / sum(B))) \quad \text{eq 3.2,}$$

where A and B are each vector encoding the number of reads for each guide in low and high sort bins respectively. Target coordinates for each sgRNAs were determined by their target PAM coordinates. Coordinates for the *GATA1* CREs are obtained using HCR-FlowFISH CASA CRE annotation (ENCFF413WYU).

##### Bootstrap sampling analysis for simulating CRISPR screens performed at various sequencing depths

Bootstrap sampling analysis for sequencing depth was performed using ENCODE standard *guide quantification* files, which record the number of sequencing reads that map to each sgRNA sequence in a given library. Each CRISPR screen comes with two guide quantification files. In case of sorting-based screen approaches (e.g. FlowFISH), one file quantifies the number of mapped sequencing reads in low-expression sorted bins (labeled as "A"), while the other file quantifies those in high-expression sorted bins (labeled as "B"). In case of growth-based screen approaches, we quantify using samples collected from an earlier time point ("A") and a later time point ("B"). To simulate an experiment with sequencing depth of *d*, we sampled with replacement total $N \times d$ number of reads independently from each A and B, where N is the number of distinct sgRNAs in a library.

For the CRISPR screens used for the bio-replicate reproducibility and dropout analyses, reads were sampled independently for each of the two bio-replicates (A1, A2, B1, B2). sgRNAs that had 0 mapped reads in any one of A1, B1, A2, and B2 were excluded from the analyses. At each *d*, 100 independent bootstrap samples were generated to be used for dropout and bio-replicate reproducibility analyses (**Fig. 3F,G**).
For the dropout simulation analysis , we defined dropout sgRNAs as those that resulted in less than 10 sampled reads from either $A_{sampled}$ or $B_{sampled}$. For bio-replicate reproducibility analysis, we computed Pearson correlation of $\log_2$FC effect sizes ($log_2((1 + A_{sampled})/(1 + B_{sampled}))$) from every pair of bootstrap samples, one coming from bio-replicate 1 and the other coming from bio-replicate 2.

**Peak caller comparisons**

*aggrDESeq2*

For each experiment, read counts of individual sgRNAs for the initial and final time points were obtained from the guideQuant files. Differential abundance testing was performed using the DESeq2 package with default parameters, with contrasts defined such that the average $\log_2$FC of sgRNAs more abundant in the final time point or high-expressing bin have positive values. Next, 100bp bins were tiled across chromosomes containing perturbations. Coordinates for individual sgRNAs were adjusted based on the perturbation modality (Cas9 cutting: ±10 bp of PAM, dCas9: ±10 bp of PAM, dCas9-KRAB: ±150 bp of PAM) and intersected with the bins. For every 100bp bin, a significance value was calculated using Fisher's method for aggregating p-values with the unadjusted DESeq2 p-values as input. The aggregated p-values were then FDR-adjusted. Significant bins were defined as FDR<0.01. Note that sgRNAs that intersect >1 bin contribute to the calculations for all overlapping bins. This was repeated without filtering out sgRNAs with GuideScan specificity scores<0.2.

*CASA*

sgRNA guideQuant files were parsed to provide genomic mapping coordinates of the protospacer sequence and raw guide counts per experimental condition in the CASA input format. We ran a containerized deployment (https://hub.docker.com/r/sjgosai/casa-kit; version 0.2.3) on the Google Cloud Platform using a wrapper script provided in the CASA GitHub repository (https://github.com/sjgosai/casa). CASA was run using a sliding window of 100 bp width and step size and a ROPE threshold of 0.693 (i.e., the default settings). As in previous work[5], peaks which are supported by at least 10 sgRNAs and shared between two bio-replicates are reported.

*CRISPR-SURF*

sgRNA guideQuant files were parsed according to the input format required for CRISPR-SURF (in particular, converting PAM coordinates to protospacer coordinates). Then, `SURF_count` was run with the options `-nuclease cas9 -pert crispri` to produce an input file for the deconvolution. `SURF_deconvolution` was run using the `-pert crispri` option, and the resulting negative_significant_regions.bed was used to identify positive regulators of expression with FDR<0.05. `CRISPR_SURF` was run using the provided Docker container using Singularity.

*MAGeCK*

sgRNA guideQuant files, coordinate expansion were performed similarly as above. 100 bp bins were created by taking the first, most upstream, coordinate position among all sgRNAs in the respective screening library, then creating 100 bp bins until reaching the most downstream sgRNA coordinate position. Expanded coordinate sgRNAs were then intersected with the bins. MAGeCK was run using the default parameters (`--norm-method=median -sort-criteria=negative -remove-zero=none -gene-lfc-method=median`), and only the significance values corresponding to the expected effect size direction for each screen (negative for the growth screens, positive for the FlowFISH screens) were used to calculate significance, which was calculated similarly as above.

*RELICS*
The sgRNA guideQuant files were prepared to provide genomic coordinates and raw counts of each sgRNA in the standard input format for RELICS. The sgRNAs overlapping promoter regions and exons of each target gene were labeled as functional sequences for CRISPRi screens and CRISPRCas9 screens, respectively. CRISPR systems used for each screen were specified for RELICS. Then, the functional sequences were identified for each screen using the default settings for RELICS v.2.0 (`min_FS_nr:30, glmm_negativeTraining:negative_control`).

*Pairwise Jaccard similarity*
For each method, peaks were loaded and a set was constructed with all nucleotides in the tiled region called significant. For each pair of peak calling methods, the Jaccard similarity was computed as:

$$\frac{|A \cap B|}{|A \cup B|}.$$

For the "Canonical Elements", we used the coordinates of the *GATA1* promoter (hg38 chrX:48786330-48786733), eGATA1 (chrX:48782816-48783227), and eHDAC6 (chrX:48800584-48800859).

*Effect sizes within peaks*
For the comparison of the distribution of guide effects ($log_2FC$) for the sgRNAs falling within peaks identified by different peak callers we started by using eq 3.2 to calculate the $log_2FC$ for each guide. We then picked the sgRNAs that overlap with the called peaks for each analysis tool and plotted the $log_2FC$ of the filtered sgRNAs.

*Nucleotide overlap with annotations*
Peaks identified by different CRISPR cCRE callers were intersected with ENCODE (DHS: ENCSR000EKS; H3K27ac: ENCSR000AKP) and SCREEN annotations (**Supplementary Table 15**).

*Intersection of CRE calls*
Significant CRE calls from each peak caller were intersected using bedtools multiinter. The output was used to generate the upset plots using the 'upset' function within the R package 'UpSetR'.

**Comparison of timepoints**

A CRISPRi growth screen with sgRNAs tiling the *GATA1* locus (ENCSR719QWB) was used to analyze the effect of timepoint selection. CASA peak calls were generated as described above. Relatedly, a CRISPRi HCR-FlowFISH screen at the *GATA1* locus (ENCSR917XEU) was inspected for dropout due to potential growth effects.

**Strand specific quantification of sgRNA effect sizes**

All CRISPR screens used in this analysis had specific gene targets (CRISPRi growth screen tiling across *GATA1* locus, and HCR-FlowFISH), and their sgRNAs were unambiguously labeled as either template or coding strand targeting sgRNAs depending on which strand their protospacers are located relative to the transcriptional directions of their target genes (**Fig. 6A,B**). For the *GATA1* CRISPRi growth screen, sgRNAs were filtered for GuideScan-aggregated CFD specificity score >0.2 to remove sgRNAs with off-target growth effects. Then, we labeled each sgRNA as gene-targeting if its PAM sequence was located between 2,000 base pairs downstream of TSS and TES. The 2,000 spacers were used to exclude gene-body targeting sgRNAs that are TSS-proximal and affect promoter activities. sgRNAs with PAM sequences located between 2,000 base pairs upstream of the TSS and the TSS itself were labeled promoter-targeting, and all other sgRNAs were labeled as "outside" (**Fig. 6C**). Refgene annotations were used to identify TSS and TES for each gene, and for genes with multiple isoforms, isoforms with the highest levels of K562 Pol-II Chip-seq signals (ENCFF914WIS, signal P values) at both the TSS and TES were used. 3 out of 20 HCR-FlowFISH experiments were excluded for this analysis **(Fig. 6D)** as they had less than 5 tested protospacers located within template-strand promoters, coding-strand promoters, template-strand gene bodies, or coding-strand gene bodies.

##### Data availability

The data will be made available in the online ENCODE portal. sgRNA counts for the *GATA1* titration experiments are provided in **Supplementary Table 18**. Public repositories to visualize CRISPR screen data and results from **Fig. 1** and **Fig. 6** are listed below:

- Fig. 1: https://data.cyverse.org/dav-anon/iplant/home/joh27/track_hub_fig1/hub.txt
- Fig. 6: https://data.cyverse.org/dav-anon/iplant/home/ohjinwoo94/track_hub_fig6/hub.txt

##### Code availability

The code for CASA can be found at https://github.com/sjgosai/casa. The code for using GuideScan2 to design sgRNAs for all cCREs can be found at https://github.com/schmidt73/encode_pipeline. The code used for other analyses will be made available in an online repository.

##### Public datasets accessed

Accession IDs for public datasets used in this study are provided in **Supplementary Table 15**.

##### Author contributions

S.K.R., J.T., D.Y., and A.K. conceived of the study. S.K.R., D.Y., J.T., J.W.O., L.R.B., S.J.G., L.L., A.M-S., B.R.D., and X.R. analyzed data. A.M-S. performed *GATA1* HCR FlowFISH coverage titration experiments. I.G., D.Y., L.R.B., J.W.O., and Y.L. curated and designed the ENCODE CRISPR screening portal, and S.K.R., D.Y., A.M-S., J.W.O., L.R.B., J.M.E., I.G., and Y.L. developed the file formats. J.W.O. and A.M-S. generated public repositories to visualize CRISPR screen data and results. J.W.O. and L.R.B. generated public repository for all code used for analyses in the paper. I.G. wrote the tutorial for navigating screening data on the ENCODE portal. L.R.B. performed literature review for design tools and analysis methods. H.S., D.Y., J.T., J.M., C.L., and Y.P. designed the genome-wide ENCODE SCREEN cCRE sgRNA libraries. M.A.B. advised analyses. S.K.R., D.Y., J.T., J.W.O., L.R.B., S.J.G., L.L., A.M-S., B.R.D., X.R., and J.M.E. wrote the paper, with revisions from all authors. S.K.R., M.C.B., M.A.B., J.M.E., A.K., and T.E.R supervised and developed the project. M.C.B., M.A.B., W.J.G., C.A.G., A.K., T.E.R., P.C.S., and Y.S. acquired funding.

The authors would like to note that when reporting this publication, all authors have agreed that co-listed authors can be listed in any order, including arranging themselves first to best highlight the equal contribution.

**Acknowledgements**

We thank members of the ENCODE4 Consortium, in particular, the members of the ENCODE4 Functional Characterization Centers who have provided feedback throughout this project. We thank Stephanie Calluori, Eileen Cahill, Dan Gilchrist, Mike Pazin, Briana Nunez, Jessica Au, and other NHGRI staff for helping to organize ENCODE conferences, jamborees, working group meetings, and providing feedback, as well as Anna Shcherbina for setting up shared computational infrastructure during the jamborees. We thank members of the ENCODE DCC for collecting, curating, and making the ENCODE data portal accessible. We thank Brian D. Cosgrove for providing helpful suggestions.

**Conflict of Interest Statements**

A.K. is scientific co-founder of Ravel Biotechnology, is on the scientific advisory board of PatchBio, SerImmune, AINovo, TensorBio and OpenTargets, is a consultant with Illumina and owns shares in DeepGenomics, Immuni and Freenome. C.A.G. is a co-founder of Tune Therapeutics and Locus Biosciences, and an advisor to Tune Therapeutics and Sarepta Therapeutics. C.A.G. is an inventor on patents and patent applications related to CRISPR epigenome editing. J.T. and M.C.B. acknowledge an outside interest in Stylus Medicine. L.L. is currently employed by Sana Biotechnology. P.C.S is a co-founder of and consultant to Sherlock Biosciences and Board Member of Danaher Corporation. She is a shareholder in both companies. W.J.G. is a co-founder of Epinomics and an adviser to 10X Genomics, Guardant Health and Centrillion.
